# Sequence adaptations in the intracellular domain of Symbiosis receptor-like kinase (SymRK) promoted infection thread progression in root nodule primordia

**DOI:** 10.64898/2026.05.07.723153

**Authors:** Maria Spezzati, A. Isabel Seidler, Martina K. Ried-Lasi, Jonathan Jelen, Julia Buchner, Miriam Abele, Tora Fougner-Økland, Cédric Beckers, Andreas Klingl, Christina Ludwig, Katarzyna Parys, Martin Parniske

## Abstract

The uptake of nitrogen-fixing bacteria into living plant cells and the intracellular accommodation of arbuscular mycorrhiza (AM) fungi requires the plasma membrane-localised Symbiosis Receptor-like Kinase (SymRK). AM is widespread across terrestrial vascular plant lineages, while the nitrogen-fixing root nodule symbiosis (RNS) is restricted to one clade within the eurosids. This distribution led to the concept that *SymRK* was adopted during evolution to mediate RNS. Comparative analyses revealed that SymRK orthologs from the eurosid clade support RNS while SymRK from the phylogenetically distant species *Solanum lycopersicum* (tomato) does not. To dissect the molecular basis for this different functionality, we carried out complementation analyses of the *Lotus japonicus symrk-3* mutant which is unable to form AM or RNS. Domains swap chimera from the tomato and *L. japonicus* SymRK orthologs revealed that the intracellular domain of *L. japonicus* SymRK is necessary and for cortical infection thread (IT) and symbiosome development at 21 days post inoculation. Notably, this signalling specificity could be overcome by ectopic expression of tomato SymRK, pointing to altered protein dosage as a potential determinant of function. Consistent with this idea, SINA family E3 ubiquitin ligases interacted with and ubiquitinylated *L. japonicus* SymRK, but not tomato SymRK. In yeast two hybrid analysis, the interaction of SymRK with SINA2 and SINA4 depended on the C-terminal intrinsically disordered tail region of *L. japonicus* SymRK. We conclude that the SymRK intracellular domain evolved interaction capabilities with SINA E3 ligases which correlates with its ability to support RNS.

## Introduction

The majority of land plant lineages are able to form arbuscular mycorrhiza (AM) with *Glomeromycota* fungi (Schüβler et al., 2001). In this symbiosis, a network of fungal hyphae extends into the soil to forage for phosphate, water, and other nutrients. Within cortical cells of the host root, highly branched fungal structures called arbuscules are formed, which mediate the exchange of these nutrients for photosynthetically derived carbon and lipids supplied by the plant (Keymer et al., 2017; Luginbuehl et al., 2017). In addition to AM, a limited number of species within the four plant orders Fabales, Fagales, Cucurbitales and Rosales, collectively referred to as the FaFaCuRo clade, have evolved the capacity to form the nitrogen-fixing root nodule symbiosis (RNS) (Doyle et al., 1997; Kistner and Parniske, 2002; Doyle, 2011). In many legumes (Fabaeae), gram-negative rhizobia enter the root through plant-derived root hair infection threads (ITs) (Ardourel et al., 1994). Driven by the pressure exerted by dividing rhizobial cells, ITs extend toward the actively proliferating cortical cells of the developing nodule (Brewin 2000, Parniske 2018). In the central tissue of the nodule, rhizobia are released into membrane-bound compartments, where they differentiate into bacteroids, organelle-like structures specialised for nitrogen-fixation (Kereszt et al., 2011).

The intracellular accommodation of fungi and bacteria in AM and RNS, respectively, shows striking morphological similarities and is governed by a shared genetic program (Kistner and Parniske, 2002). The underlying genes, known as Common Symbiosis Genes (CSGs), are conserved across plant species and are thought to have been adopted from the more ancient genetic program required for AM development during the evolution of RNS (Kistner and Parniske, 2002). Mutations in CSGs, including the *Symbiosis Receptor-like Kinase* (*SymRK*), impair both symbioses, underscoring their central and conserved roles in symbiotic development of host root cells. A key step in the evolution of RNS was the emergence of the *Predisposition-Associated Cis-regulatory Element* (*PACE*) within the promoter region of the *NODULE INCEPTION* (*NIN*) gene (Schauser et al., 1999; Cathebras et al., 2026). Because CSGs are present across plant species that form AM, it is possible that their function in RNS required discrete sequence adaptations that support RNS. Evidence for such adaptation has previously been reported for *SymRK* (Markmann et al., 2008).

*SymRK* encodes a plasma membrane-localised receptor-like kinase containing a malectin-like domain (MLD), leucine-rich repeats (LRRs) and an active serine/threonine kinase domain (Yoshida and Parniske, 2005; Abel et al., 2024). Mutant analysis in model legumes, including *Lotus japonicus* (hereafter *L. japonicus*) have established an essential role for SymRK in both AM and RNS. In *L. japonicus symrk* (previously called *sym2*) mutants, aborted infection attempts of AM fungi in the root epidermis were observed, characterised by swollen fungal hyphae inside the lumen of dead plant cells (Bonfante et al., 2000; Stracke et al., 2002; Demchenko et al., 2004; Kistner et al., 2005). During RNS, *symrk-3* fails to form ITs in response to rhizobia and consequently does not form nitrogen-fixing root nodules (Stracke et al., 2002). Similar phenotypes were observed on roots of plants carrying the *symrk-10* allele (Markmann et al., 2008). The *symrk-10* allele encodes a SymRK version in which aspartic acid 738 (D738) in the activation loop is replaced with asparagine (N), rendering SymRK catalytically inactive (Perry et al., 2003; Yoshida and Parniske, 2005). Interestingly, the role of SymRK in cell layers below the epidermis remains to be clarified. SymRK appears dispensable for arbuscule formation (Demchenko et al., 2004, Ivanov et al., 2024). However, RNAi-mediated silencing of *SymRK* orthologs in *Sesbania* and *Medicago* impairs rhizobial release and symbiosome formation in the central tissue of the root nodule (Capoen et al., 2005; Limpens et al., 2005). These observations reveal that SymRK may exert symbiosis-relevant functions beyond the epidermis.

Trans-species complementation studies in the *L. japonicus symrk-10* mutant using *SymRK* orthologs from non-FaFaCuRo AM-forming species, further suggest that *SymRK* underwent functional adaptation to support RNS and that these changes predated the acquisition of RNS. Specifically, SymRK from tomato (*Solanum lycopersicum* (euasterids I Solanales) restored AM but not RNS. Curiously, even a SymRK variant from *Tropaeolum* (Brassicales), far outside the FaFaCuRo clade but within the eurosid II clade, restored both root endosymbiosis (Markmann et al., 2008). Because all orthologous genes were expressed under the native *L. japonicus SymRK* promoter, these functional differences are likely due to sequence polymorphisms in the SymRK proteins. However, the specific protein-level polymorphisms and their functional consequences that underpin the adoption of SymRK for bacterial root endosymbiosis remain unresolved. In this work, we address this question through a comparative analysis of SymRK orthologs from *L. japonicus,* which forms both AM and RNS, and tomato, which forms AM but not RNS.

## Results

### Tomato SymRK restores development of root nodule primordia-like lateral organs on *Lotus japonicus symrk-3* roots, but the progress of infection threads is impaired

We compared the ability of *L. japonicus SymRK* (*Lotus*; *LjSymRK*) and its ortholog from *Solanum lycopersicum* (tomato; *SlSymRK*) to restore AM or RNS in the *L. japonicus symrk-3* mutant. Hairy root transformed *L. japonicus symrk-3* plants expressing tomato or *Lotus SymRK* under the control of the 4.9 kb native *LjSymRK* promoter are hereafter referred to as *SlSymRK/symrk-3* and *LjSymRK/symrk-3,* respectively. Both *LjSymRK* and *SlSymRK* restored the passage of AM fungi through the epidermis in the *symrk-3* mutant (Fig. S1). In contrast, the two constructs differed strikingly in their ability to support RNS development.

#### Infection thread development

To compare the early infection phenotypes in *LjSymRK/symrk-3* and *SlSymRK*/*symrk-3* roots, the number of root hairs with deformations, entrapments of bacteria with root hair curling and infection threads (ITs) were quantified 7 days post inoculation (dpi) with *Mesorhizobium loti (M.loti) Ds*Red (Fig. 1A). Empty vector (EV)/*symrk-3* was used as negative control. Among responsive root hairs on *SlSymRK/symrk-3*, 13% formed ITs, compared with 33% on *LjSymRK*/*symrk-3* roots (Fig. 1A). This difference was not statistically significant. However, significantly more *LjSymRK/symrk-3* root hairs responded with curling and entrapments of bacteria (52% vs 26%) compared to *SlSymRK/symrk-3* (Fig. 1A). An opposite trend was observed for root hairs with deformations (61% vs 15%) (Fig. 1A). The significantly higher ratio of root hair deformations on *SlSymRK/symrk-3* suggest a compromised IT initiation, but this is uncoupled from IT progression once formed. We conclude that tomato S*lSymRK* can support IT formation in root hairs of *L. japonicus symrk-3* mutant roots.

**Figure 1.**
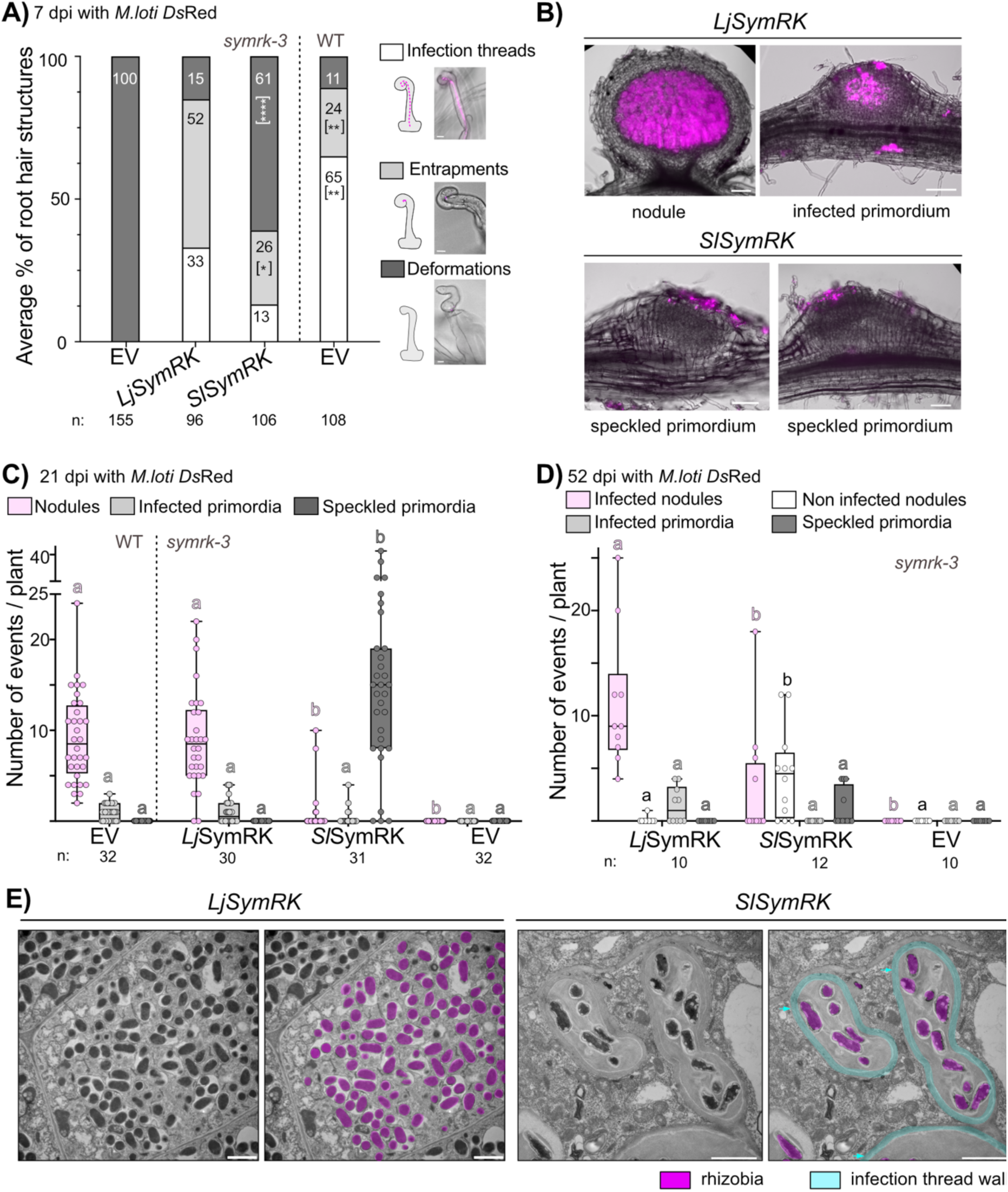
The progression of infection threads into the central tissue is blocked on *L.japonicus symrk-3* roots expressing *SlSymRK*. A) Average percentages of infection threads, entrapment events and deformed root hairs per plant on transformed roots using the same genetic constructs as described in B. Representative microscopy images of the different root hair structures formed in response to *M. loti Ds*Red inoculation are shown on the left side of the graph. Merged bright field and *Ds*Red channel images are shown. Note that infection threads formed when *symrk-3* was complemented with *Sl*SymRK but with a lower frequency as compared to *Lj*SymRK. Statistical significance was assessed using two-way ANOVA followed by Šídák’s multiple comparison test. *Sl*SymRK and EV/WT were compared against *Lj*SymRK. Asterisks indicate significance levels: * p = 0.05 – 0.01, ** p = 0.01 – 0.005, *** p < 0.005. For each SymRK construct, 3 to 5 plants were selected for quantitative analysis. Scale bars: 20 µm. n: total number of root hairs counted. B) Representative microscopic images of longitudinal sections of nodule organogenesis events observed on roots of *symrk-3* mutant transformed with *L. japonicus (Lj)* (nodule, infected primordium) or tomato *(Sl)* (speckled primordia) *SymRK* under the native *LjSymRK* promoter. Merged bright field and *Ds*Red channel (magenta) images are shown. Note that speckled primordia refer to primordia-like structures with fluorescent signal only in the outer cell layers but not in the central tissue. Speckled primordia were exclusively observed on *SlSymRK*/*symrk-3*. Phenotypes were assessed at 21 days post-inoculation (dpi) with *M. loti Ds*Red. Scale bar: 100 µm. C) Boxplot displaying the number of nodules, primordia and swellings scored on either wild-type Gifu (WT) roots transformed with the Empty Vector (EV), or on *symrk-3* roots transformed with *Lj*SymRK, *Sl*SymRK or the EV. Statistical significance was assessed using two-way ANOVA followed by Dunnett’s multiple comparison test. P-value < 0.05 between groups is indicated with different letters. n: number of transformed plants analysed. D) Hairy roots of *L. japonicus symrk-3* transformed with Empty Vector (EV), *LjSymRK*, *SlSymRK* all expressed under the control of the endogenous *SymRK* promoter were incubated for 52 dpi with *M. loti Ds*Red. Infected and non-infected nodules were distinguished by the presence of fluorescent *Ds*Red signal. Note the increased number of non-infected nodules on *Sl*SymRK-transformed roots. Statistical significance was assessed using two-way ANOVA followed by Tukey’s multiple comparison test for the different categories. P-value < 0.05 is indicated with different letters. n: number of transformed plants analysed. E) Electron micrographs of cross sections of a nodule (*Lj*SymRK/*symrk-3* roots) and a speckled primordium (*Sl*SymRK/*symrk-3* roots) 21 dpi with *M.loti Ds*Red. Rhizobia are highlighted in pink, while infection thread walls in cyan. Note that in the section of a speckled primordium, bacteria are surrounded by a membrane and imbedded in a matrix, surrounded by plant cell wall material (cyan arrowheads), resembling an infection thread structure. Multiple infection thread (IT) sections are observed within one cell, suggesting IT curling or branching, but neither bacterial release nor symbiosome formation could be detected. Scale bar: 1 µm. Experiment was repeated three times independently.

#### Organogenesis

Nodule organogenesis on *LjSymRK/symrk-3* and *SlSymRK/symrk-3* hairy roots was analysed 21 dpi with *M. loti Ds*Red. Both constructs supported the development of root nodule primordia, but they differed markedly in morphology and infection progress. Infection status was quantified by the presence of a fluorescent signal from *M. loti Ds*Red. On *Lj*SymRK/*symrk-3* as well as on the positive control Empty Vector (EV)/Gifu wild type, mature infected nodules formed at 21 dpi with *M. loti Ds*Red (Fig. 1B, C). In contrast, the formation of *“*speckled” primordia was detected on 90% of *Sl*SymRK/*symrk-3* root systems investigated (Fig. 1B, C, Fig. S2). We called them “speckled” because the central tissue of these primordia-like structures was not colonised by *M. loti Ds*Red. Instead, fluorescence was consistently observed at the surface of these primordia-like structures in top-view images and in cells above the central tissue in cross-sections (Fig. 1B, Fig S2). Together with the formation of ITs in *Sl*SymRK/*symrk-3* roots (Fig. 1A) these observations suggest that ITs failed to progress into the inner tissue by 21 dpi. Microscopic inspection of longitudinal cross-sections revealed that a large, non-quantified, subset of these speckled primordia was characterized by pronounced elongation along the root axis. This elongation was not observed in infected nodule primordia on *Lj*SymRK/*symrk-3* roots (Fig. 1B, Fig. S2). To test the stability of the transition block into the central tissue in *SlSymRK/symrk3*, the restoration of RNS development was inspected after extended incubation times. After 52 dpi, infected nodules were present on all root systems of *LjSymRK/symrk-3,* and on 33% of *SlSymRK/symrk-3* roots (Fig. 1D). Notably, we observed that non-infected nodules, defined by the lack of a fluorescent signal from *M. loti Ds*Red, formed on 75% of *SlSymRK/symrk-3* root systems, whereas only a single non-infected nodule was found on one *LjSymRK*/*symrk-3* root system (Fig. 1D). Moreover, speckled primordia were observed exclusively on *SlSymRK/symrk-3* roots at 52 dpi. Compared with 21 dpi (Fig.1C), their frequency was reduced, occurring in 33% rather than 90% of the total root systems (Fig. 1D). The observed increase in the number of non-infected nodules on *SlSymRK/symrk3* roots over time – combined with a reduced number of speckled primordia – suggests that nodule organogenesis is restored, while the infection block is stable in the majority of nodules even over an extended incubation time.

The progression of ITs into deeper cell layers of nodules and speckled primordia at 21 dpi was investigated by electron microscopy (Fig. 1E). As expected, densely packed symbiosomes containing bacteroids were detected in the central tissue of the nodule from *LjSymRK/symrk-3* roots. In speckled primordia (only found on *LjSymRK/symrk-3* roots), the meristematic central tissue was free of bacteria, and symbiosomes were not detected (Fig. 1E, Fig S3). In cell layers underneath the epidermal tissue, bacteria were present inside plant cells but clearly separated from the plant cell cytoplasm by a cell wall-like structure, forming a bacteria-enclosing compartment. Within the lumen of this compartment the bacteria were embedded in a matrix material that differs from the cell wall material by lower electron density (Fig. 1E, Fig S3). We consider these to be IT-like structures given their resemblance to images published previously (Jordan et al., 1963; Dixon, 1964). Since such structures accumulated in the outer cell layers of speckled primordia but were absent from the central tissue, we conclude that *SlSymRK/symrk-3* supports IT progression into outer layers but is impaired in promoting further progression or spreading into the nodule central tissue.

### Sequence polymorphisms in the intracellular domain of SymRK are critical for infection progression into the central tissue

Tomato and *Lotus* SymRK orthologs exhibit substantial amino acid sequence divergence (53% identity) and structural domain variation exemplified by one fewer Leucine-rich repeat in tomato variant (Markmann et al., 2008). To pinpoint sequence polymorphisms functionally linked to infection progression into the central nodule tissue, we designed a series of chimeric *SymRK* variants for the expression of domain swap proteins. Guided by sequence conservation, the three domains of *Lj*SymRK and *Sl*SymRK – the extracellular domain (ECD), transmembrane region (TM) and intracellular domain (ICD) – were exchanged between the orthologs (Fig. 2). All chimeric constructs were expressed under the control of 4.9 kb endogenous *LjSymRK* promoter.

**Figure 2.**
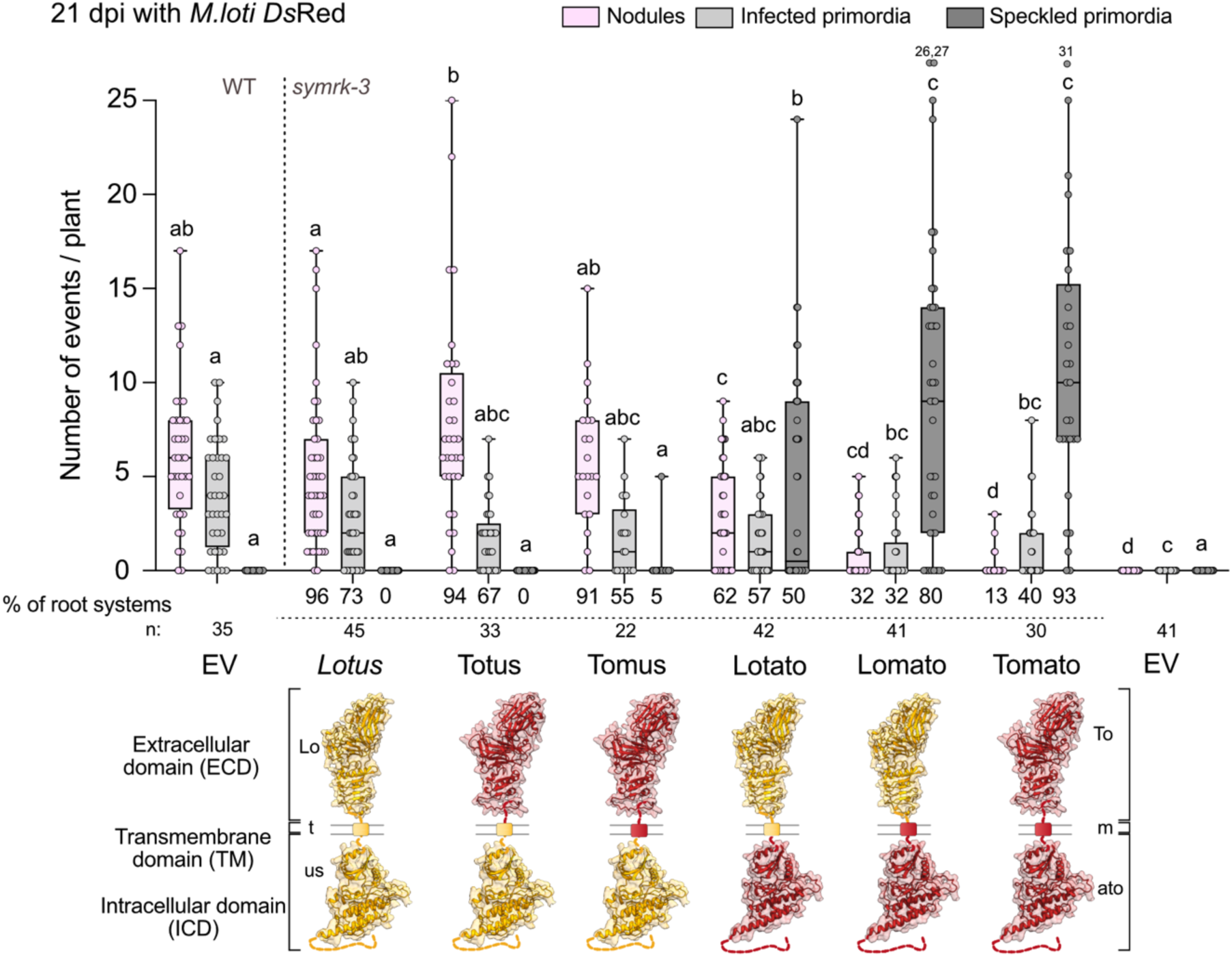
The intracellular domain of *Lj*SymRK facilitates rhizobia progression into the cortex. Number of infected nodules, infected primordia and speckled primordia observed on roots of *L. japonicus* Gifu (WT) transformed with an empty vector (EV) as a positive control, or of *symrk-3* expressing the *Lotus* or tomato orthologs of *SymRK* or the indicated domain swap constructs. All constructs were driven by the *Lotus SymRK* promoter. Chimeras containing *Lotus* ICD *e.g* Totus and Tomus, fully restored the nodule and infected primordia numbers to the level observed with *Lotus* SymRK. Note that speckled primordia were observed on roots of *symrk-3* transformed with Lotato and Lomato constructs, which contain the tomato ICD. The *symrk-3* mutant transformed with EV did not develop any organogenesis structure. The percentage of transformed root systems where nodules, infected primordia or speckled primordia were observed is indicated below the box plot. Statistical significance was assessed by two-way ANOVA followed by Tukey’s multiple comparison test for the different categories. P-value < 0.05 is indicated with different letters. n: number of transformed plants analysed. The 3D structures of SymRK *Lotus* and tomato were predicted using Alphafold3.

Two of these chimeric variants, called Lomato and Totus, were able to restore wild type-like epidermal passage of the AM fungus, validating that the encoded chimeric proteins were expressed and functional (Fig. S1). We tested RNS competence at 21 dpi with *M. loti* and observed mature nodules on more than 90% of root systems of the positive controls (*EV*/Gifu wild type and *LjSymRK/symrk-3*) (Fig. 2). Similar nodule frequencies were restored in *symrk-3* roots transformed with the chimeric constructs Totus and Tomus, which both share a common feature: they contain the ICD of *Lj*SymRK. In contrast, when *symrk-3* roots were transformed with constructs encoding chimera carrying the tomato ICD, *e.g.* Lomato and Lotato, speckled primordia were observed (Fig. 2). The only exception was a single Tomus root system, however, this construct predominantly induced nodules and infected primordia formation.

Remarkably, the proportion of root systems forming speckled primordia versus nodules in Lotato and Lomato-transformed roots is influenced by the identity of the TM region. Overall, the presence of *Lj*SymRK TM (Lotato vs Lomato) leads to significantly more infected nodules (62% vs 32% of root systems) and significantly fewer speckled primordia (50% vs 80% of root systems). Taken together, sequence features within the intracellular domain of *Lj*SymRK appear sufficient to support infection progression and colonisation of the nodule central tissue, whereas the *Lotus* SymRK transmembrane domain exerts an additional – and, at least in Lotato, ICD – independent, positive effect on RNS development.

### Tomato SymRK is able to complement the kinase dead *symrk-10* mutant

To gain mechanistic insights into the compromised functionality of tomato SymRK ICD, we investigated whether the infection block can be overcome either by increasing SymRK dosage or by leveraging the partial functionality of *Lotus* SymRK. For this, we took advantage of the *symrk-10* mutant allele, in which SymRK is expressed but in its kinase-inactive form (Yoshida and Parniske, 2005; Abel et al., 2024). We generated transgenic hairy roots of *symrk-10* expressing *LjSymRK*, *SlSymRK* and two selected chimeric constructs Tomus and Lotato (Fig. S5).

Except *EV/symrk-10,* which was used as a negative control, all constructs tested could rescue the *symrk-10* mutant phenotype 21 dpi with *M. loti Ds*Red to a level similar to that reached on *LjSymRK/symrk-10* roots (Fig. S5 A). Thus, the additive effect of expressing the kinase-dead *Lj*SymRK in the *symrk-10* mutant together with *Sl*SymRK is sufficient to restore infection progression into the central nodule tissue. This ability of *Lj*SymRK^D738N^ encoded by the *symrk-10* mutant can be explained in at least three, not mutually exclusive, scenarios: (i) increased SymRK protein dosage, (ii) provisioning of an interaction surface for key signalling intermediates, and/or (iii) *Sl*SymRK trans-phosphorylating the kinase domain of *Lj*SymRK^D738N^ variant rescuing the specific phospho-code required for functional symbiotic signal transduction (Fig. S5 B).

### *Lotus* and tomato SymRK intracellular domains display distinct autophosphorylation patterns

We compared the *in vitro* auto-phosphorylation sites of *Lotus* and tomato SymRK-ICD expressed *in E. coli*, purified by affinity chromatography followed by size-exclusion chromatography and subjected to *in vitro* kinase assays. The kinase-dead version *Lj*SymRK^K622E^ served as negative control (Supp. Table 3). By mass spectrometry we identified multiple putative auto-phosphorylation sites in the ICD of SymRK from *Lotus* and tomato that were not detected with confidence on the negative control *Lj*SymRK^K622E^ indicating that both kinases were purified in their catalytically active forms (Fig. S6). Although we cannot fully exclude the possibility that the low confidence sites (1-2/3 replicates) that were found on the kinase dead version were phosphorylated by kinases of *E. coli* (Supp. Table 3).

Tomato and *Lotus* SymRK-ICDs exhibited notable differences in both the number and positions of the phosphorylated serine (S) and threonine (T) residues (Fig. S6). Overall, 12 phosphorylation sites were detected in tomato SymRK ICD, each detected at least four times across six replicates. In comparison, 35 phosphorylation sites were detected with high confidence in *Lotus* SymRK ICD, with a 42% overlap with phosphorylation sites reported by Abel et al. (2024)(Fig. S6). Inspection of the AlphaFold 3 structural models showed that the sites were surface-exposed and distributed along the entire ICD. In the lower lobe and the unstructured C-terminal tail (lower lobe from amino acid 747 onward), a substantially higher number of sites was observed in *Lj*SymRK as compared to *Sl*SymRK. We categorized the modified S and T residues as conserved or divergent based on sequence alignment between the two SymRK proteins. The conserved S751 and S754 on *Lj*SymRK (S732 and S735 on *Sl*SymRK) in the activation segment were phosphorylated in both ICDs. Yet, to our surprise, many of the conserved sites, present in both proteins, were phosphorylated differently, with *Lj*SymRK carrying more sites. Nine S/T were conserved and phosphorylated in both proteins; 16 S/T were conserved but only phosphorylated in *Lotus* but not in tomato. We also found tomato– and *Lotus*-specific sites (9 S/T in *Lotus* vs 2 S/T in tomato), revealing divergent phosphorylation patterns between these two receptor proteins. Collectively, these results highlight differences between *Lotus* and tomato SymRK in both amino acid sequence and distinct phosphorylation patterns of the intracellular domain.

### Ectopic expression of tomato *SymRK* promotes IT progression and RNS development in *symrk-3*

Ectopic expression of *LjSymRK* or *Arachis* kinase domain induces spontaneous nodule organogenesis in the absence of rhizobia (Ried et al., 2014; Saha et al., 2014). To test if ectopic expression of tomato *SymRK* can bypass the infection block in nodule central tissue, we selected three promoters to drive the expression of *Lotus* and tomato *SymRK*. We compared the endogenous promoter used in the previous experiments (Fig. 1 and 2), the *LjUbiquitin* promoter and additionally an endogenous promoter with the chimeric transcription activator VP16-Gal4 (*proSym^enhanced^*) that leads to increased transcript levels of genes driven by promoters equipped with this module (Fig S7 A). Here we used this module with the aim to increase transcript – and thus conceptually SymRK protein – abundance.

Hairy roots of the *symrk-3* mutant transformed with different *promoter:SymRK* constructs were analysed for spontaneous nodule formation in the absence of *M. loti*, and RNS phenotypes 21 dpi with *M.loti Ds*Red. None of the *promoter:SlSymRK* variants led to spontaneous nodule organogenesis (Fig. S7 B). In contrast, spontaneous nodule organogenesis occurred on all *proUbi:LjSymRK/symrk-3* root systems, confirming previous results (Ried et al., 2014). Moreover, two root systems (15%) transformed with the enhancer construct *proSym^enhanced^:LjSymRK* exhibited spontaneous nodule organogenesis (Fig. S7 B).

Upon inoculation with *M. loti Ds*Red, we observed infected nodules and primordia on all *EV*/Gifu wild type, and on most of the transformed roots of *proSymRK:LjSymRK/symrk-3* (90%) and *proSym^enhanced^:LjSymRK/symrk-3* (90%). Compared to *proSymRK:LjSymRK /symrk-3*, fewer infected nodules were found on average on *proUbi:LjSymRK/symrk-3*, and most root systems (57 %) formed non-infected nodules (Fig. 3).

**Figure 3.**
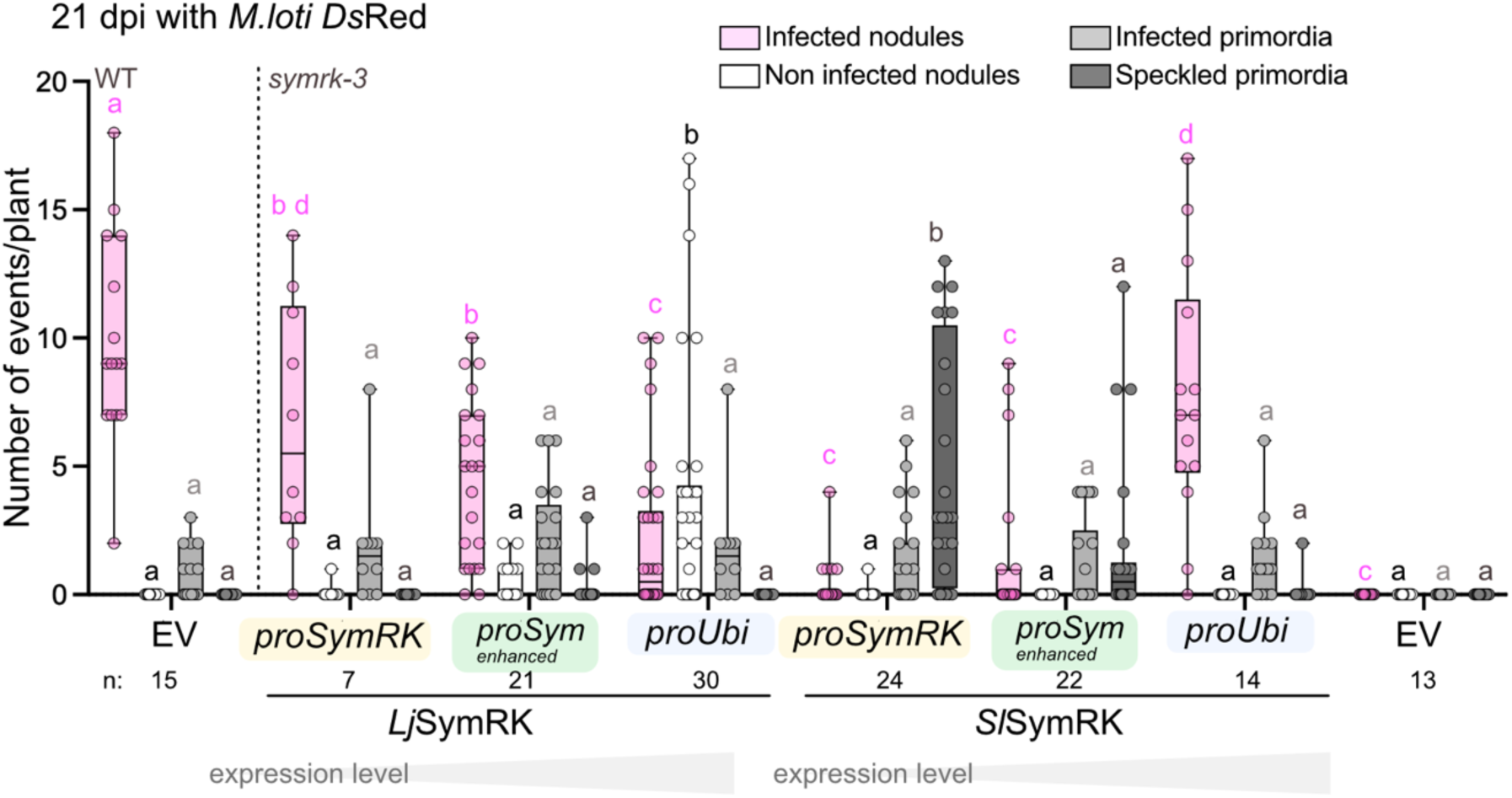
*SlSymRK* driven by the *Ubiquitin* promoter restored root nodule symbiosis in *symrk-3*. Tomato and *Lotus SymRK* expressed under different promoters as described in Supplementary Figure 7A, were used for complementation of *symrk-3* using hairy root transformation. EV in *L. japonicus* Gifu (WT) and *symrk-3* was used as positive and negative control, respectively. Infected and non-infected nodules, primordia and swellings were counted 21 dpi with *M. loti Ds*Red. Note the different phenotypes caused by *SlSymRK* expression driven by the *proSym*^enhanced^ and the *Ubiquitin* promoter (*proUbi*) compared to the SymRK endogenous promoter (*proSymRK*). A significantly lower number of speckled primordia was observed for both *proSym*^enhanced^ and *proUbi,* while significant increase of infected nodules was only observed for *proUbi* compared to the native *proSymRK.* No non-infected nodules were found on any of the *Sl*SymRK transformed roots, while *proUbi:Lj*SymRK*/symrk-3* roots carried a higher number of non-infected than infected nodules. Statistical significance was assessed by two-way ANOVA followed by Tukey’s multiple comparison test for the different categories. P-value < 0.05 is indicated with different letters. n: number of transformed plants analysed.

Strikingly, *Sl*SymRK expressed under the control of *proUbi* complemented *symrk-3* mutant to a similar level as *Lj*SymRK under its native promoter, with 92% of root systems carried infected nodules (Fig. 3). *SlSymRK* expressed from *proSym^enhanced^* only partially restored infected nodule formation, inducing infected nodules on 27% of root systems. Nonetheless, the numbers of speckled primordia were significantly reduced compared to the endogenous promoter (Fig 3). This suggests that the promoter used for *SlSymRK* expression has a significant effect on its capacity to support IT progression into the root nodule central tissue of the *symrk-3* mutant.

### *Lotus* and tomato SymRK intracellular domains differ in their interactions with, and ubiquitinylation by, E3 ligases

Based on the observation that *Sl*SymRK driven by native *SymRK* promoter can restore infected nodules in *symrk-10* mutant (Fig. S5) and driven by a *proUbi* can complement *symrk-3* mutant (Fig. 3), we hypothesise that receptor dosage could be important for SymRK function. Receptor stability is often regulated by ubiquitinylation, a post-translational modification dependent on the activity of E3 ubiquitin ligases (Rodriguez-Furlan et al., 2019).

Multiple studies have identified E3 ubiquitin ligases that interact with the ICD of SymRK, including members of the SEVEN IN ABSENTIA (SINA) family and PUB1 (Plant U-box E3 ligase 1) (Den Herder et al., 2012; Vernié et al., 2016). These findings collectively support a model in which PUB1 and SINA4 affect SymRK stability.

A semi-quantitative yeast two-hybrid assay (Y2H) was employed to test the interaction between intracellular domains of *Lotus* and tomato SymRK and SINA E3 ligases (Den Herder et al., 2012), as well as the *L. japonicus* ortholog of *Medicago truncatula* PUB1 (Vernié et al., 2016). A version of PUB1 carrying a deletion of the N-terminal domain (hereafter PUB1^ΔN^) was used for all the assays in this manuscript. For interaction studies, a mutated version in the U-box domain (PUB^ΔN, W319A^) was used, which was found not to affect interaction with SymRK (Vernié et al., 2016). We confirmed the interaction between SINA1, SINA2, SINA3, SINA4 and PUB1^ΔN^ proteins with *Lotus* SymRK-ICD (Fig. 4A) in line with previous reports (Den Herder et al. 2012; Vernié et al. 2016). Importantly, tomato SymRK ICD did not interact with any of the tested SINA family members, but did interact with PUB1^ΔN, W319A^ (Fig. 4B).

**Figure 4.**
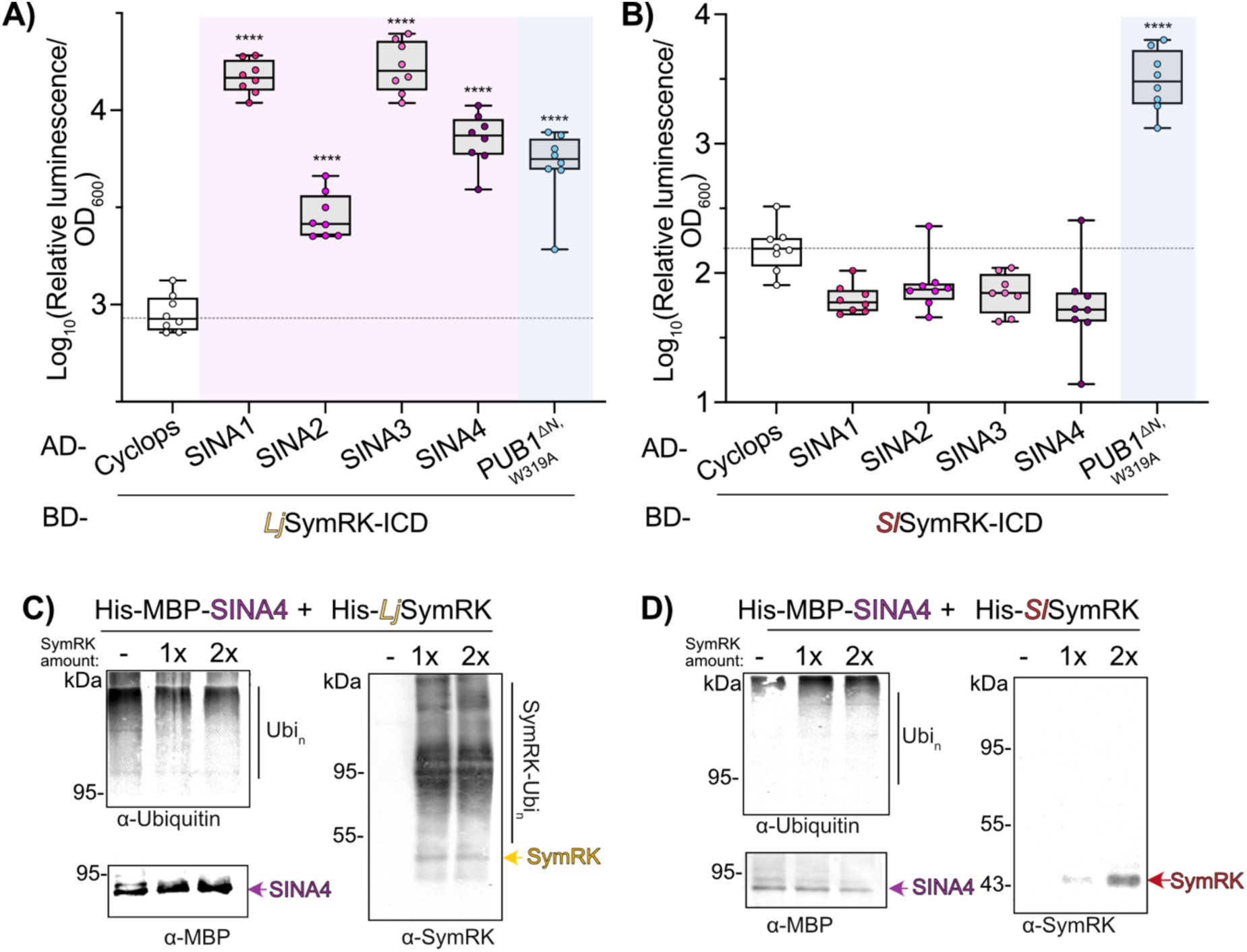
The intracellular domain of *Lj*SymRK specifically interacts with SINA E3 ligases and is ubiquitinylated by SINA4. A) and B) Yeast two-hybrid interaction studies between *Lotus* (A) and tomato (B) SymRK-ICD and E3 ubiquitin ligases PUB1^ΔN^ and SINA1-4. The boxplot represents normalized luciferase signals measured for independent yeast colonies co-transformed with constructs encoding the SymRK ICD fused to the Gal4 Binding domain (BD) and SINA1, 2, 3 or 4 or PUB1 fused to the Gal 4 Activation Domain (AD). Cyclops-AD was used as a negative control. The significant luminescence signal intensity for *Lj*SymRK ICD suggests interaction with both SINA E3 ligases and PUB1^ΔN, W319A^, whereas the luminescence signal for *Sl*SymRK suggests interaction only with PUB1^ΔN, W319A^. A yeast strain carrying an integrated *proGal2:Luciferase* reporter was used. Statistical significance was evaluated by one-way ANOVA followed by Dunnett’s multiple comparison test. Asterisks indicate p-values < 0.05. C) *In vitro* ubiquitinylation assay with His-MBP-SINA4 and His-*Lj*SymRK ICD proteins purified from *E. coli*. The ubiquitinylation patterns of both proteins were analysed through Western Blot with anti-Ubiquitin, anti-SymRK and anti-MBP antibodies. 1x and 2x amounts of SymRK were used, as indicated above the membranes. Note the presence of high-molecular weight signal for the membrane incubated with anti-SymRK antibodies. The yellow arrowhead marks non-modified SymRK ICD (43 kDa), while the purple arrowhead marks SINA4 (80 kDa). D) In a setup similar to C), His-TRX-*Sl*SymRK ICD was used as a substrate in an *in vitro* ubiquitinylation assay with His-MBP-SINA4. Note the absence of high molecular weight signal for the membrane incubated with anti-SymRK antibodies. The red arrowhead marks non-modified His-TRX-*Sl*SymRK (60 kDa), while the purple arrowhead marks SINA4 (80 kDa).

We further tested the differential interaction between SymRK orthologs and the inactive versions of SINA2, SINA4 and PUB1^ΔN^ with Fluorescence Lifetime Imaging Microscopy-Förster Resonance Energy Transfer (FLIM-FRET) (Singh et al., 2014). For this, the intracellular domain of *Lotus* or tomato SymRK was fused to nuclear-localization signal (NLS) and mGFP fluorescent protein, while NLS and mCherry fluorescent protein were fused to the N-terminus of PUB1^ΔN, W319A^, SINA2^C64S^ and SINA4^C66S^ (Fig S9). The only exception was the *Sl*SymRK ICD – PUB1^ΔN, W319A^ interaction, where a different orientation of fluorescent proteins was used (mCherry for SlSymRK and mGFP for PUB1^ΔN, W319A^). We co-expressed fusion constructs encoding the donor (mEGFP) and acceptor (mCherry) proteins in *Nicotiana benthamiana* mesophyll leaves and subsequently measured the reduction in fluorescent lifetime of the donor. Our results confirm that PUB1^ΔN, W319A^ interacts with both SymRKs, while SINA2^C64S^ and SINA4^C66S^ interact exclusively with *Lotus* SymRK (Fig. S11), in agreement with the Y2H data (Fig. 4).

To determine whether the differential interaction between SymRK and SINA4 is linked to sequence polymorphism between tomato and *Lotus* SymRK ICDs, or to sequence differences between the SINA orthologs, we repeated Y2H interaction studies using tomato putative orthologs of SINA4 and PUB1 identified by OrthoFinder analysis. Hereafter, we refer to these as *Sl*SINA4 and *Sl*PUB1^ΔN^, respectively (Fig. S11). Interestingly, we found that *Sl*SINA4 can interact with *Lj*SymRK ICD but not with *Sl*SymRK ICD in our Y2H assay (Fig. S12 A). In contrast, we observed interaction of *Sl*PUB1^ΔN^ with *Sl*SymRK (Fig. S12 B). Thus, while the interaction between PUB1 and SymRK ICD is conserved across species, the differential interaction between tomato and *Lotus* SymRK with SINA4 is likely a result of a sequence polymorphism in the SymRK binding surface.

### *Lj*SymRK, but not *Sl*SymRK, is ubiquitinylated by SINA4

To test whether SymRK interaction with SINA2, SINA4 and PUB1^ΔN^ leads to ubiquitinylation of SymRK, we performed *in vitro* ubiquitinylation assays. Ubiquitinylation of tomato and *Lotus* SymRK-ICD by SINA2, SINA4 and PUB1^ΔN^ was analysed via western blots with SymRK-specific antibodies (Fig. 4C, D and S10). We detected a high-molecular-weight ladder pattern corresponding to poly-ubiquitinylated *Lj*SymRK, but not for *Sl*SymRK, in the presence of SINA4 (Fig. 4C, D) but not of ubiquitinylation SINA2 (Fig. S10A). No self-ubiquitinylation of SINA2 was detected in the absence of SymRK (Fig. S10). The lack of self– and trans-ubiquitinylation activity for SINA2 suggests that, under our experimental conditions, SINA2 does not function as an active E3 ligase. In contrast, PUB1^ΔN^ ubiquitin ligase ubiquitinylated the ICDs of both *Lj*SymRK and *Sl*SymRK (Fig. S10B, C).

These patterns could be confirmed through LC–MS analysis of lysine residues modified by ubiquitinylation (Fig. S13 and S14). 12 lysine residues on *Lj*SymRK ubiquitinylated by SINA4 were detected in all the three replicates; in contrast, no modified lysine residues were detected on *Sl*SymRK with high confidence. (Fig. S14). Interestingly, no ubiquitinylation sites were detected on *Lotus* SymRK lysine residues residing in the terminal alpha-helix and unstructured C-tail. For PUB1^ΔN^, we detected ubiquitinylation of 16 lysines in *Lj*SymRK (79% of all lysine residues in the ICD) and 13 in *Sl*SymRK (72% of all lysine residues in the ICD), distributed across the entire ICD model (Fig. S14). This suggests high promiscuity of PUB1^ΔN^ towards its substrates. In summary, *Lj*SymRK interacts with both SINA2 and SINA4, while *Sl*SymRK does not interact with either, even not with the tomato SINA orthologues. *In vitro* ubiquitinylation could only be detected by SINA4 but not with SINA2. On the other hand, PUB1 interacts with and ubiquitinylates both SymRK orthologs, *Lotus* and tomato. These *in vitro* findings demonstrate that the ubiquitinylation pattern of SymRK depends on the interacting E3 ligase and, consequently, that its protein homeostasis depends on interactions with diverse E3 ligases.

### The polymorphic C-terminal tail of SymRK determines the interaction capability with SINA4 and SINA2

We hypothesised that the differential interaction between SINA2 and SINA4 and *Lotus* versus tomato SymRK results from amino acid polymorphisms within the SymRK ICD. To narrow down ICD regions relevant for the interaction, we engineered chimeric proteins by reciprocal domain swaps. Specifically, the kinase domain (K) of SymRK from *Lotus* or tomato was fused to either the juxtamembrane (J) region or the C-terminal tail (C-tail; C) of different SymRK^JKC^ orthologs (Fig. 5), and tested for interaction with SINA2 and SINA4 in a Y2H assay. We found that the swap of the *Lotus* (L) C-tail with the tomato (T) C-tail (ICD^LLT^) abolished interaction with SINA4 and SINA2, whereas the reciprocal swap (ICD^TTL^) enabled this interaction (Fig. 5). Similar results were obtained using deletion constructs of either juxtamembrane (*LjSymRK*^ΔJ^), C-terminal tail (*LjSymRK*^ΔC^) or both subdomains *LjSymRK*^ΔJC^ (Fig. S17). Only the full-length *LjSymRK* ICD and the *LjSymRK*^ΔJ^ variant, both of which retain an intact *Lotus* C-tail, maintained interaction with SINA4 (Fig. S15). Together, these results support the conclusion that the C-terminal tail of *Lotus* SymRK is critical for its interaction with SINA4 and SINA2.

**Figure 5.**
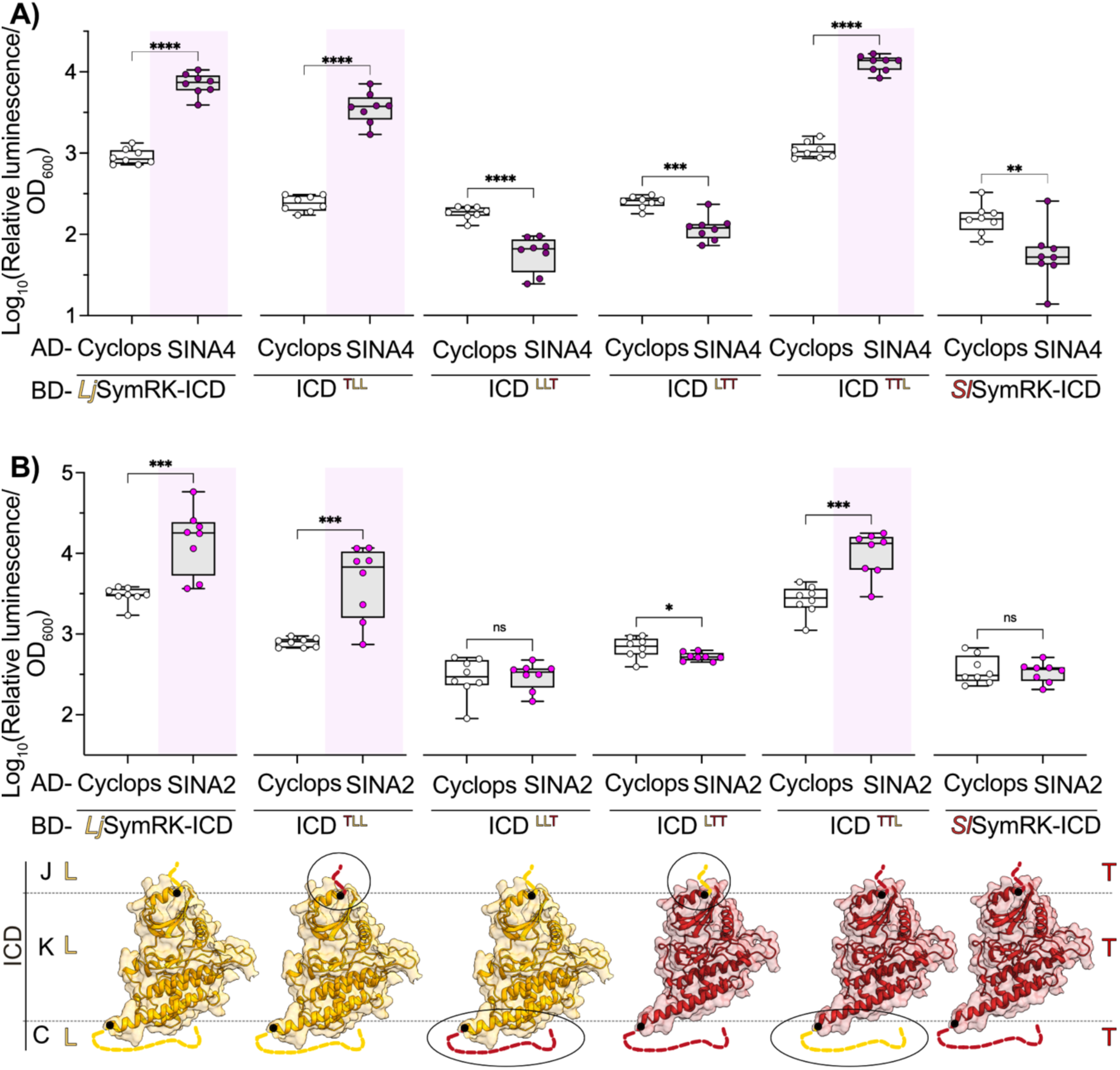
The C-terminal region of *Lotus* SymRK determines its interaction with SINA2 and SINA4. A) and B) Yeast two-hybrid interaction studies between SINA4 (A) or SINA2 (B) and different variants of the SymRK ICD. The boxplots represent normalized luciferase signal measured for independent yeast colonies co-transformed with a construct encoding the SymRK ICD fused to the Gal4 Binding domain (BD) and SINA4 or SINA2 fused to the Gal 4 Activation Domain (AD), respectively. Cyclops-AD was used as a negative control. A series of SymRK ICD chimeric variants was tested by swapping the juxtamembrane region (J), kinase domain (K) or the unstructured C-tail (C) between *Lotus* and tomato, as illustrated in the Alphafold3 models below the graph. Note that the luminescence signal intensity for *Lj*SymRK ICD, ICD^TLL^ and ICD^TTL^ indicates their interaction with both SINA2 and SINA4. On the other hand, the interaction with both SINAs is lost when C-tail of *Lotus* is swapped for C-tail from tomato (ICD^LLT^). A yeast strain carrying an integrated *proGal2:Luciferase* reporter was used in the studies. Statistical significance was evaluated by one-way ANOVA followed by Dunnett’s multiple comparison test. Asterisks indicate p-values < 0.05.

We examined sequence polymorphisms within the C-tail of selected SymRK orthologs (Fig. S4). We hypothesized that, if the SINA2 and SINA4 interaction is required for infection thread progression to the central nodule – a process blocked in *SlSymRK/symrk-3* – then the key functional sites for this interaction in the C-tail should be conserved in legumes, and possibly at least as far as the complementation capable *Tropeolum* (Markmann et al., 2008). Based on a sequence alignment of the unstructured region of the C-tail of *Lotus* and tomato SymRK orthologs, we observed low amino acid sequence identity of around 43% (Fig. 4). By comparing sequence logos of Eurosid orthologs to non-Eurosid angiosperms in this region, we observed signatures that are enriched in Eurosid SymRK orthologs. In particular, two motifs were enriched in Eurosid orthologs relative to other angiosperms (Fig. S4). These could underlie differential post-translational regulation mechanisms or interactions important for signalling.

## Discussion

### SymRK orthologs differ in their ability to restore root nodule symbiosis

In this study, we performed functional complementation experiments to assess and compare the ability of tomato SymRK, derived from a species that forms AM but not RNS, with that of *L. japonicus SymRK* to support RNS. On *SlSymRK/symrk-3* roots, we observed the development of speckled primordia, at very high numbers. The structures resemble nodule primordia in which nodule organogenesis proceeds normally, but infection threads fail or are significantly delayed to progress into the nodule’s central tissue (Fig. 1, Fig. S2, S3). Consistent with this interpretation, after prolonged incubation, most nodules remain non-colonised (Fig. 1D). These results support the hypothesis proposed previously that, sometimes in its evolutionary ancestry potentially at the root of the eurosids, SymRK underwent functional adaptations that were necessary and formed a prerequisite for its role in RNS (Markmann et al., 2008).

Interestingly, phenotypes related to the observed speckled primordia induced by tomato SymRK have been described previously for SymRK variants defective in MLD release, in which rhizobia accumulate at the apex of primordium-like structures in *L. japonicus* and *Medicago truncatula* (Antolin-Llovera et al., 2014; Li et al., 2018; Chakrabarti et al., 2024). Because MLD release destabilises the receptor and may thereby influence protein abundance (Antolin-Llovera et al., 2014), these observations, together with our findings, point to receptor homeostasis as an important determinant of infection thread progression. We therefore propose that changes in SymRK stability, and consequently in its interaction dynamics within specific nodule cell layers, contribute to coordinating the transition of the infection thread into the central tissue. Consistent with the reduced requirement for *SymRK* in cortical arbuscule formation (Demchenko et al., 2004; Ivanov and Harrison, 2024), our findings suggest that SymRK function in RNS is subject to a strict, tissue-specific regulation. We found that *Sl*SymRK driven by the native *Lj*SymRK promoter fully rescued the kinase-dead *symrk-10* mutant (Fig. S5). SymRK homodimerization has recently been postulated (Chakrabarti et al., 2024). In light of these results, we hypothesise that *Sl*SymRK forms receptor complexes with the catalytically inactive *Lj*SymRK^D738N^, thereby generating phosphorylation patterns on the kinase-dead ICD of *Lj*SymRK^D738N^, including in the critical alpha-I motif (Abel et al., 2024), that are required for RNS signalling. Alternatively, partial functionality of *Lj*SymRK^D738N^ could be related to the attraction of intracellular interacting proteins or increased dosage of SymRK, which together contribute to the compromised functionality of *Sl*SymRK. In light of our overexpression results, increased receptor dosage appears to be a parsimonious explanation.

The functional complementation results in *symrk-10* mutants feature interesting differences compared to previous published assay. Markmann et al. (2008) did not observe nodule primordia formation or nodule organogenesis induction when introducing tomato *SymRK* into a *Lotus symrk-10* mutant background. For the experiments described by Markmann et al. (2008) the plants were grown under different temperature and light regimes. For example, the setting used traditional tubular fluorescent lamps yielding lower light intensities and a distinct spectral range. Such differences in cultivation conditions are a possible cause of the discrepancies in phenotypic readouts reported in the two studies.

### SymRK overexpression can bypass receptor specificity constraints during RNS

Our data show that ectopic expression of *Sl*SymRK can fully complement RNS in the presence of *M. loti* (Fig. 3). By using an enhanced version of the *SymRK* promoter, we found that increased expression of tomato SymRK reduces the number of empty elongated primordia (Fig. 3). This result is in line with previous research where other orthologs of SymRK, such as rice, could also complement *symrk* mutant when driven by a *LjUbiquitin* promoter (Li et al., 2018). These results indicate that receptor specificity can be overridden when protein levels are excessively high, likely outcompeting negative regulatory components and/or overriding limits otherwise imposed by intrinsic affinity constant values of protein-protein interactions.

Consistent with this, hypermorphic phenotypes resulting from receptor overexpression have been reported previously (Wang et al., 2001). In the absence of rhizobia, ectopic expression of *Lj*SymRK under a *LjUbiquitin* promoter in *L. japonicus symrk-3* (Ried et al., 2014) or expression of the *Arachis* SymRK ICD driven by *35S* promoter (Saha et al., 2014), leads to spontaneous nodule organogenesis (Fig. 3) and activation of AM transcriptional marker genes (Ried et al., 2014).

### E3 ligases as modulators of SymRK signalling and homeostasis

SINA-family E3 ligases interact with *Lj*SymRK (den herder and Fig. 4). Based on the single-cell RNA-sequencing database of *L. japonicus* roots (Frank et al., 2023), we found that *SINA2* and *SINA4* are co-expressed with *SymRK* in more than 23% of root cells, whereas *SINA1* and *SINA3* are co-expressed in less than 9% of root cells at 10 dpi with *M. loti* (Fig. S8). This prompted us to investigate the biochemical properties of the SINA2 and SINA4 proteins, although it is likely that the SINA1 and SINA3 proteins also play a role in SymRK homeostasis regulation.

*Lj*SymRK can interact with tomato ortholog of SINA4, while *Sl*SymRK cannot. These results suggest a gain or loss of SymRK’s interaction capability over evolutionary timescales. SINA4 may act as a regulator of *Lj*SymRK homeostasis; its ectopic expression leads to SymRK degradation and relocalisation when overexpressed in *N. benthamiana* leaf cells (Den Herder et al., 2012). Interestingly, the negative regulatory function of SINA4 seems to be functionally relevant for infection thread formation in RNS, but no phenotype was observed for AM (Den Herder et al., 2012). The abnormal cortical ITs found in stable lines over-expressing SINA4 (Den Herder et al., 2012) also suggest an abnormal release of rhizobia in the nodule.

These findings support a model in which SINA4 acts in a spatially or temporally restricted manner to modulate SymRK abundance during infection progression. Following this hypothesis, *Sl*SymRK lacking the interaction with SINA4 in the deeper cell layers may lead to mis-regulation of receptor dynamics: without proper degradation or relocalisation, SymRK accumulates in a non-functional or aberrant state, contributing to the block in infection progression. Additionally, multiple members of SINA family did not interact with *Sl*SymRK, so we cannot exclude a function for the other SINAs in regulating SymRK stability.

Unexpectedly, SINA2 did not show self-ubiquitinylation activity *in vitro*, despite conservation of its catalytic RING domain. This suggests that SINA2 may not function as a canonical E3 ligase in this context. One possibility is that SINA2 requires a specific E2 conjugase, which we did not test *in vitro*. Alternatively, SINA2 may function as a “decoy” E3 ligase binding to SymRK, without promoting ubiquitinylation. In such a scenario, SINA2 could modulate receptor homeostasis or signalling output through competitive binding rather than degradation, representing a potentially distinct mode of RLK regulation.

As SINA4 appears to be a specific regulator of *Lj*SymRK function in RNS, PUB1 may provide a more general, conserved layer of control important for the establishment of AM and RNS. PUBs are often described as regulators of RLKs (Mbengue et al., 2010; Lu et al., 2011; Stegmann et al., 2012; Vernié et al., 2016; Liu et al., 2018). Consistently, PUB1 appears to function as a conserved E3 ligase, as it interacts with and ubiquitinylates both *Lj*SymRK and *Sl*SymRK (Fig. 4, Fig. S12). This idea is indirectly supported by the phenotype observed in *Medicago pub1* mutant where both AM and RNS are affected (Vernié et al., 2016). While we observe *in vitro* ubiquitinylation, we currently lack evidence for PUB1-mediated degradation of SymRK. This raises the intriguing possibility that PUB1 may regulate SymRK through non-degradative ubiquitinylation (e.g., K63-linked chains), influencing signalling complex assembly, localisation, or activity. Another hypothesis is that PUB1-mediated degradation of SymRK, so far tested only in *N. benthamiana,* requires either a symbiotic trigger or a specific SymRK phosphorylation state.

### Sequence polymorphisms in the C-terminal tail and divergent phosphorylation patterns likely contribute to functional diversification of SymRK

We identified the unstructured C-terminal tail as a key determinant of the interaction with the SINA2 and SINA4 E3 ubiquitin ligases (Fig. 5). Notably, the C-terminal tail of RLKs frequently serves as a regulatory domain. The C-tail of Arabidopsis ERECTA receptor acts as an autoinhibitory domain with its activity is switched on or off via phosphorylation-dependent recruitment of either PUB30/31 or BRI1 KINASE INHIBITOR (BKI1) (Chen et al., 2025). Similarly, BRASSINOSTEROID INSENSITIVE 1 (BRI1) C-tail fine-tunes the kinase activity, serving as a negative regulatory domain (Wang et al., 2021). These examples highlight a widespread mechanism in which the C-tail maintains RLK homeostasis and enables precise responses to external stimuli (Wei et al., 2025).

In addition to sequence polymorphisms, post-translational modifications – such as phosphorylation – can influence molecular interactions and downstream signalling outputs. We found that tomato and *L. japonicus* ICDs differ in their phosphorylation patterns, with *L. japonicus SymRK* showing a higher degree of auto-phosphorylation at S/T sites (Fig. S7). This suggests that the functional divergence between orthologs may also be due to the presence or absence of specific phosphorylation codes. For example, we did not detect phosphorylation of the gatekeeper tyrosine (Y670 in *A. hypogaea* SymRK), which has been reported to be essential for the epidermal-to-cortex infection transition in RNS (Saha et al., 2016). Moreover, despite the conservation of these residues in the tomato SymRK ortholog, the alpha-I motif (Abel et al., 2024) is only partially phosphorylated. A phospho-mimetic mutation in the alpha-I motif of SymRK induces spontaneous nodule formation (Abel et al., 2024), indicating that an auto-active conformation of SymRK-receptor complexes can bypass normal signal discrimination and promote ligand-independent activation of the symbiotic signal pathway. These results suggest that the missing phosphorylation sites possibly contribute to *Sl*SymRK’s complementation capacity.

In summary, our data support a model in which the evolutionary adoption of SymRK for RNS required sequence features of its intracellular domain related to receptor homeostasis. Homeostasis (and potentially phosphorylation patterns of its ICD) may be the key for SymRK signalling capacity necessary for infection thread progression in the nodule central tissue. This conclusion is based on three main pillars, 1) the differential SINA interaction mediated by C-tail polymorphisms, 2) the ability to override the specific features by overexpression, and 3) the ability of the *symrk-10* allele encoding a kinase-dead yet structurally intact SymRK version, to enable the *SlSymRK* version to restore RNS. Especially the last point opens the possibility that the observed LjSymRK-specific phosphorylation patterns may be critical for its function in RNS. Together, these findings provide a mechanistic framework of features SymRK requires to fulfil its RNS-specific role while retaining its ancestral function in AM.

## Materials and Methods

### Biological material

*Lotus japonicus* genotypes used: Gifu B-129 (wild type, *symrk-3*, *symrk-10*) (Handberg and Stougaard, 1992; Stracke et al., 2002; Perry et al., 2003). For AM experiments plants were inoculated with *Rhizophagus irregularis* DAOM 197198 (Symplant GmbH & Co. KG, Darmstad Germany) using a chive nurse plant system (*Allium schoenoprasum*). RNS phenotypes were scored after inoculation with *Mesorhizobium loti* strain MAFF303099 expressing the *Discosoma* sp. red fluorescent protein *Ds*Red. (Maekawa et al. 2008). *Agrobacterium rhizogenes* strain AR1193 was used for hairy root transformation (Offringa et al., 1986). For transient transformation of *N.benthamiana*, *Agrobacterium tumefaciens* GV3101 was used*. E.coli* TOP10 strain was used for the cloning of synthetic plasmids. *E.coli* Rosetta pLaqI (Novagen) strain (SymRK ICDs) and *E. coli* BL21 (SINA2, SINA4, PUB1) were used for protein expression. Yeast strain Y1880 with stably integrated Firefly *Luciferase* gene was used for Yeast-Two Hybrid assays. To drive the expression of *Luciferase*, the *proGal2* from *S. cerevisiae* strain AH109 was used.

### Plasmid construction

All plasmid constructs were generated using the Golden Gate strategy (Engler et al., 2008; Binder et al., 2014) with adaptions to the cut-ligation protocol as described in (Chiasson et al., 2019). All primers used and plasmids generated are listed in Supplementary tables 1 and 2. Primers and plasmids were designed using CLC Main Workbench 20.0.4. The enhancer construct VP16-Gal4 is based on Sevin-Pujol et al. (2017).

### Plant growth conditions and hairy root transformation

*Lotus japonicus* seeds were scarified and surface-sterilized as described in Gossmann et al., 2012. Seeds were germinated on ½ Gamborg’s B5 medium solidified with 0.8 % Bacto™ agar on square plates (12 x 12 x 1.7 cm) (Gamborg et al., 1968). Seeds were kept in dark for three days before transferring to light (160 µmol m-2·s-1) and grown in long-day conditions (16 h/8 h light/dark) in a Polyklima growth cabinet at 24 °C. Six-days-old seedlings were subjected to hairy root transformation as described before (Charpentier et al., 2008). Transformed roots were selected using a fluorescence stereo microscope Leica, M165FC based on positive NLS-GFP signal.

### Inoculation for symbiosis phenotyping

#### M. loti inoculation

One week before rhizobia inoculation, plants with transformed hairy roots were moved to a nitrogen-reduced FAB medium (500 µM MgSO_4_·7H_2_O; 250 µM KH_2_PO_4_; 250 µM KCl; 250 µM CaCl_2_·2H_2_O; 100 µM KNO_3_; 25 µM Fe-EDDHA; 50 µM H_3_BO_3_; 25 µM MnSO_4_·H_2_O; 10 µM ZnSO_4_·7H_2_O; 0.5 µM Na_2_MoO_4_·2xH_2_O; 0.2 µM CuSO_4_·5H_2_O; 0.2 µM CoCl_2_·6H_2_O; pH 5.7) solidified with 0.8% Bacto™ agar in square plates. Plants were inoculated in Weck^®^ jars (SKU 745 or 743) containing 300 ml of sand-vermiculite mixture (2:1) and 30 ml of nitrogen-reduced FAB medium containing *M. loti Ds*Red adjusted to a final optical density at 600 nm (OD_600_) of 0.05. Plants were grown for the following three weeks in long day conditions (light intensity: 275 µmol·m-2·s-1) in a growth chamber at 24 °C. For spontaneous nodule analysis, similar procedure was performed, excluding inoculation with *M. loti Ds*Red.

#### AM inoculation

Chive *(Allium schoenoprasum*) roots colonised with AMF *R. irregularis* were used for inoculation of *Lotus* roots. Surface sterilized chive seeds were sown in sand-vermiculite and around 5000 spores of *R. irregularis* DAOM 197198 were added in 1⁄4 Hoagland medium (Hoagland, 1938). Colonized chive roots were subsequently used as inoculum for mycorrhized chive propagation. Chive roots grown at least 8 weeks in pots with AM, were cut in ±1 cm pieces and mixed with sand:vermiculite mixture (2:1). *L. japonicus* hairy roots were transferred to 200 ml of the mixture containing colonized chopped chive roots in 8×7 cm pots. Each pot was watered with 20 ml of autoclaved FAB medium low phosphate three times per week.

### Phenotypic analysis and quantification of infection events

#### Quantification of RNS structures

*M. loti* – inoculated plants were harvested and roots were cleaned from sand and vermiculite in water using a fine brush. The shoots were cut off, and the roots were fixed in 4% formaldehyde in 50 mM PIPES (pH 7) by vacuum infiltration. After fixation, roots were washed three times with PIPES, pH 7, before stored in 50mM PIPES, pH 7 at 4 °C. The different nodulation phenotypes were assessed using an upright epifluorescence microscope (Leica DM6 B). Infected and non-infected nodules were distinguished by the presence or absence of a *Ds*Red (representing *M.loti Ds*Red) signal inside the nodules. Primordia and swellings were discriminated by the presence of the *Ds*Red signal in the central part of the primordia or accumulated in the outer cell layers, respectively. The formation of entrapments and infection threads was screened using *Ds*Red signal for identification of *M. loti* using an upright epifluorescence microscope (Leica DM6 B).

#### Sections of nodules, primordia and lateral organs

Nodules and primordia plus 2 mm of the surrounding root were embedded in 6% low-melting agarose. Transversal, 50 μm thick sections were consecutively cut off with a vibrating-blade microtome (Leica VT1000 S), and inspected for *Ds*Red signal using a Leica DM6 B epifluorescence microscope.

#### AM phenotyping

For AM phenotyping, the roots were thoroughly cleaned from sand and vermiculite using tap water and a fine brush. Non-transformed roots, identified by the lack of GFP fluorescence of the transformation marker, were removed by cutting them off under a stereo microscope (Leica, M165FC). AM fungal structures were stained by acetic acid ink staining as described previously (Vierheilig et al., 1998). In brief, roots were harvested in 10% KOH, incubated at 90 C for 15 min, and incubated in 10 % acetic acid for 10 min. Afterwards, roots were stained with 5% black ink in 5% acetic acid and distained in 5% acetic acid. Root colonization was inspected using a bright field microscope.

### Ultrastructure analysis via Electron Microscopy

For chemical fixation, root samples with nodules or primordia were immersed for 4 weeks in 2.5 % glutaraldehyde in 75 mM cacodylate buffer with 2 mM MgCl_2_ pH 7.0 to initiate the fixation process. Samples were washed with buffer three times and then post-fixed with 1 % osmium tetroxide (OsO₄) 2h20 minutes to crosslink unsaturated fatty acids and enhance membrane contrast. After repeated rinsing with distilled water, the samples underwent dehydration through a graded acetone series (10–100 %), with 2 % uranyl acetate included in the 20 % acetone step to provide additional contrast. Epoxy resin infiltration was carried out using a graded series of resin-acetone mixtures, culminating in a final incubation in 100 % Spurr resin. After polymerized at 60 °C for 24 h, the samples were trimmed and 60 nm thin sections were obtained using an ultramicrotome. These sections were placed on coated grids and post-stained with 3% lead citrate for 2 minutes prior to transmission electron microscopy at 80 kV in the zero-loss mode using a Zeiss EM912 (Zeiss, Oberkochen, Germany) equipped with a 2k x 2k slow-scan CCD camera (TRS – Tröndle Restlichtverstärker Systeme, Moorenweis, Germany).

### Yeast-two Hybrid interaction studies

For yeast transformation, around 50 ml liquid YPAD medium (Supp. Table 3) was inoculated with yeast grown on YPAD plates and grown over night at 28°C, under continuous shaking at 180 rpm. The next day the OD_600_ was diluted at an OD_600_ of 0.15 in 50 ml YPAD-containing flask. Yeast cells were grown in a 200 ml Chicane flask at 28°C for 4 h, under continuous shaking at 180 rpm. Cells were pelleted at 4000 g, for 5 min. The pellet was washed with sterile dH_2_O and centrifuged again. Pellet was resuspended in 1 ml ddH2O and transferred to a 1.5 ml reaction tube. Cells were pelleted at 13000 g, for 30 s. Supernatant was removed and pellet was resuspended in 0.5 ml ddH2O. For each transformation, 50 μl of cells were aliquoted into 1.5 ml reaction tubes, containing 200 ng per plasmid (400 ng total amount for double transformation). 326 μl of cold transformation mix (36.8 % PEGG 3550, 110 mM Lithium Acetate, 0.3 mg/mL salmon sperm DNA) was added followed by immediate vortexing. Cells were incubated at RT for 15 min, followed by an incubation of 30 min at 42°C. Cells were pelleted at 13000 g for 30 s. Supernatant was removed and cells were resuspended in 500 μl of autoclaved dH_2_O. 100 μl were plated onto synthetic dropout (SD) medium lacking Leucine and Tryptophan. Cells were grown at 28°C for 3 days before performing luciferase assay. Individual colonies were inoculated in single wells of a sterile 96 deepwell plate, containing 300 μl sterile SD medium per well. Plate was closed and placed in an incubator overnight (28°C, 350 rpm). 100 μl were transferred to a transparent 96 well Sarstedt plate and OD_600_ was determined using a TECAN plate reader (Absorbance at 600 nm, 25 flashes, shake for 3s at 4 mm linear amplitude). Measured values were multiplied by a scaling factor of 12.1. Scaling factor was determined empirically by comparing OD600 of an overnight culture measured with a 1 ml cuvette in a photospectrometer, with the OD600 of 100 μl of the same culture measured in a TECAN plate reader.

Based on the obtained OD_600_ values, cultures were inoculated in a fresh 96 deep-well plate at an OD of 0.5 and grown at 28°C, 350 rpm. After 4 h, 100 μl were transferred to a transparent 96 well Sarstedt plate and OD_600_ was determined like above. From the same cultures, 100 μl were transferred to a white 96 well COSTAR plate and used for luminescence measurement. 100 μl luciferin substrate (1 mM D-luciferin in 0.1 M sodium tri-citrate, pH=3) was injected at following parameters: 3 s shaking at 3 mm linear amplitude, injection speed 200 μl/s, refill speed 200 μl/s. Luminescence values were normalised to OD600 values.

### Protein expression and purification from *E.coli*

BL21 or Rosetta expression strains were transformed with one of expression plasmids listed in Supplementary Table 2 using heat shock protocol (Froger and Hall, 2007). Single colony was select to inoculate a liquid pre-culture and grown overnight at 37°C. The following day, expression culture was initiated in final volume 500 mL-2L of LB medium with 2% of the pre-culture and incubated at 37°C until OD_600_ reached 0.4-0.6. Protein expression was induced by adding IPTG to the final concentration 0.4 mM-1mM. The flasks were shifted to 20°C and grown overnight. The next day, the cultures were pelleted by centrifugation 10.000g for 20 min at 4°C and washed once with water. If not immediately processed, the pellet was stored at –20°C.

For protein extraction, pellet was resuspended in Lysis buffer (50 mM Tris-HCl pH 7.5, 5% glycerol, 1 mM DTT, 1 mM PMSF, 100 mM NaCl, 5 mM MgCl2, 1μg/mL DNAse and 300 μg/mL lysozyme) and mechanically lysed using High Pressure Cell Disruptor. After two rounds of centrifugation at 10000g 4C, the lysate was incubated with equilibrated Sepharose beads (Ni Sepharose^TM^ Excel for SymRK ICDs, Amylose Resin NEB E8021S for SINA4 and PUB1, Glutathione-Sepharose^TM^ 4B for SINA2). The lysate was incubated for two hours with the beads rotating at 4C. Afterwards, flowthrough, washes and elutions were collected from a gravity flow column. Elution samples were concentrated using Amicon centrifugal filters. Aliquots from pellet, flowthrough, lysate, washes and elutions were mixed with Laemmli Sample buffer, incubated at 65 C for 5 minutes and visualised on a SDS gel stained with Comassie Brilliant Blue (CBB).

Concentrated elutions were loaded on a AKTA column (Superdex 200, Cytiva) for size-exclusion chromatography. Samples were collected according to the chromatogram, loaded on SDS gel and analysed after staining the gel with CBB. The elution fractions containing the proteins of interest were concentrated with Amicon filters, mixed with final 5% glycerol concentrations and stored at –70 C for further use.

### Ubiquitinylation and Phosphorylation assay *in vitro*

Ubiquitinylation assays were prepared following the the protocol from Den Herder et al., 2012. In brief: purified proteins were incubated in the 2mM MgCl2, 50mM Tris pH 7.5, 2mM ATP, 0.5 mM DTT for 2h at 30 C in the presence of E1, E2 enzymes and Ubiquitin. SymRKs were added 1:2 with the respective E3 ligases.

Reactions were analysed by SDS-PAGE followed by immunoblot analysis using anti-Ubiquitin antibodies (1/3000; Santa Cruz Biotechnology), anti-GST (1/5000, Sigma), anti-MBP (1/10000 NEB), anti-SymRK (1/2500 Agrisera) or sent for mass spectrometry-based proteomics to identify phosphorylation and ubiquitinylation ubiquitinylation sites.

### Mass spectrometry-based proteomics

#### Sample preparation

Each in vitro ubiquitinylation assay was mixed 3:1 (v:v) with 4x LDS sample buffer (NuPAGE™ ThermoFisher Scientific) and heated for 10 minutes to 70°C. In-gel trypsin digestion was performed according to standard procedures (Shevchenko et al., 2006). Briefly, 30 µl per sample was run on a Nu-PAGE™ 4%–12% Bis-Tris protein gel (ThermoFisher Scientific) for about 1 cm. Subsequently, the still not size-separated single protein band per sample was cut out, reduced (10 mM dithiothreitol in 5 mM TEAB), alkylated (55 mM chloroacetamide in 5 mM TEAB) and digested overnight with trypsin (Trypsin Gold, mass spectrometry grade, Promega). The peptides obtained were dried to completeness and resuspended in 12 μl of 0.1% formic acid in HPLC grade water. Finally, 5 μL of sample were injected per mass spectrometric measurement.

#### Mass spectrometric data acquisition

Peptides were analysed on a VanquishTM Neo UHPLC (microflow configuration; Thermo Fisher Scientific, MA, USA) coupled to an Orbitrap ExplorisTM 480 mass spectrometer (Thermo Fisher Scientific, MA, USA). Peptides were applied onto a commercially available Acclaim PepMap 100 C18 column (2 μm particle size, 1 mm ID × 150 mm, 100 Å pore size; Thermo Fisher Scientific, MA, USA) and separated on a stepped gradient from 3% to 31% solvent B (0.1% FA, 3% DMSO in ACN) in solvent A (0.1% FA, 3% DMSO in HPLC grade water) over 60 min. A flow rate of 50 μl/min was applied. The mass spectrometer was operated in DDA and positive ionization mode. MS1 full scans (360 – 1300 m/z) were acquired with a resolution of 60,000, a normalized automatic gain control target value of 100%, and a maximum injection time of 50 ms. Peptide precursor selection for fragmentation was carried out using a cycle time of 1.2 seconds. Only precursors with charge states from two to six were selected, and dynamic exclusion of 30 s was enabled. Peptide fragmentation was performed using higher energy collision-induced dissociation and a normalized collision energy of 28%. The precursor isolation window width of the quadrupole was set to 1.1 m/z. MS2 spectra were acquired with a resolution of 15,000, a fixed first mass of 100 m/z, a normalized automatic gain control target value of 100%, and a maximum injection time of 40 ms.

#### Mass spectrometric data analysis

Peptide identification and quantification was performed using the software MaxQuant (version 1.6.3.4)(Tyanova et al., 2016). MS2 spectra were searched against the Uniprot *E.coli* protein database (UP000002032, downloaded May 2024), appended with the protein sequences of *Lj*SymRK, *Sl*SymRK, SINA4 and PUB1, as well as supplemented with common contaminants (built-in option in MaxQuant). Trypsin/P was specified as proteolytic enzyme. Carbamidomethylated cysteine was set as fixed modification. Oxidation of methionine, acetylation at the protein N-terminus, phosphorylation of serine, threonine and tyrosine as well as ubiquitinylation of lysine (GG motif) were specified as variable modifications. Results were adjusted to 1% false discovery rate on peptide spectrum match (PSM) level and protein level employing a target-decoy approach using reversed protein sequences. The minimal peptide length was defined as 7 amino acids and the “match-between-runs” functionality was enabled (matching time window 0.7 min, alignment time window 20 min). Detected phosphorylation or ubiquitinylation sites on the SymRK target protein were filtered according to their reproducible detection over six (phosphorylation) or three (ubiquitinylation) technical replicates. For the phosphorylation analysis the resulting MaxQuant result tables have been deposited to the ProteomeXchange Consortium via the PRIDE partner repository. For the ubiquitinylation analysis an additional targeted data analysis strategy using the software tool Skyline (version Skyline-daily (64-bit) 26.1.1.058) (MacLean et al., 2010) has been performed. For that, the DDA data was loaded into Skyline and for all MaxQuant detected ubiquitinylation sites the respective MS1 peaks across all samples were manually reviewed and validated. The resulting Skyline documents have been deposited to Panorama Public (Sharma et al. 2018).

### Transient expression in *Nicotiana benthamiana* leaves and FLIM-FRET

*Agrobacterium tumefaciens* strain GV3101 was transformed with the expression plasmids described in Table 2. After growing them on selection plates containing Kan, Rif and Spec, one colony was used to inoculate a 5 mL liquid culture. Bacteria were grown for 24h at 28°C 180 rpm. The day after, the culture was centrifuged at RT 1500g and washed with sterile water. Bacteria pellet was resuspended in infiltration buffer (10 mM MgCl_2_). After adjusting OD_600_ of 0.2 (E3 ligases) or 0.4 (SymRKs-ICDs) the different cultures were mixed 1:1:1 with p19 and Acetosyringone was added to a final concentration of 150 µM. The tubes were incubated for 1-2h at RT in the dark and then infiltrated in the leaves of 3-4 weeks old *N.benthamiana.* After 72h, corks were collected and used for FLIM-FRET analysis. Fluorescence lifetime was measured on a TCS SP5 confocal laser-scanning microscope (CLSM) equipped with a HCX IRAPO L 25×/0.95 W objective lens (Leica, Wetzlar, Germany). GFP was excited with a Ti:Sapphire multiphoton laser (Spectra Physics, Santa Clara, U.S.) running at 80 MHz with 1.2 ps pulse lengths at 900 nm wavelength. Photons were recorded with a FLIM PMT type detector in a wavelength range of 500 to 550 nm. Signals were recorded with the TCSPC system and the SPCM photon counting software version 9.83 (Becker & Hickl, Berlin, Germany) with the FIFO imaging mode at a spatial resolution of 512×512 pixels. Lifetime calculation was performed with the SPCImage software (Becker & Hickl, Berlin, Germany). For determination of lifetime, a region of interest was set around the nucleus, pixels were binned by a factor 2 and a single exponential decay model with the Maximum Likelihood Estimation (MLE) method was applied to distinguish background and GFP signal. Scatter was fixed to zero. In the curve fitting the automatically determined internal response function (IRF) was applied.

### Phylogenetic analysis

Orthologs of SymRK, SINA4 and PUB1 were identified across 46 plant genomes using OrthoFinder v2 (Emms and Kelly, 2019). Additional SymRK orthologs for analysis of sequence polymorphisms in the C-terminal tail were identified by reciprocal bidirectional best hit BLAST against selected genomes as well as a protein RLK database generated by Ngou et al. (2024). For SINA4 and PUB1 trees, sequences were filtered to include homologs *from Lotus japonicus, Solanum lycopersicum, Medicago truncatula, Arabidopsis thaliana and Marchantia paleacea* before alignment using Clustal Omega. Trees were built using RaxML-NG (Kozlov et al., 2019) and visualised in iTOL(Letunic and Bork, 2024).

To obtain alignments of the C-tail of SymRK, sequences were aligned using MAFFT (Katoh et al., 2002). The alignment was trimmed using *Lj*SymRK as an anchoring sequence to define the starting position of the C-tail. Following trimming, sequences were realigned using MAFFT and sequences lacking a C-tail were removed. The alignment was visualised using Jalview. For sequence logos, the alignment was filtered to keep sites with >90% occupancy, and the logo was visualized using WebLogo 3 (Crooks et al., 2004). Motif enrichment analysis was done using STREME (Bailey, 2021) from the MEME suite version 5.5.9.

LjSINA1:LotjaGi1g1v0792700, LjSINA2: LotjaGi1g1v0278800, LjSINA3: LotjaGi3g1v 0320300, LjSINA4:LotjaGi1g1v0285800, LjPUB1:LotjaGi1g1v0075000, SlSINA4: Solyc03g083270.3, SlPUB1: Solyc02g072080.1, LjSymRK: LotjaGi2g1v0330500, SlSymRK: NP_001234869.1

Gene co-expression analysis was conducted using the single-cell database available online https://lotussinglecell.shinyapps.io/lotus_japonicus_single-cell/ using the 10 dpi dataset (Frank et al., 2023).

Protein modeling of *Lj*SymRK ICD and *Sl*SymRK ICD was done using Alphafold3 (Abramson et al., 2024) and using the ChimeraX software 1.6.1(Meng et al., 2023).

### Data visualization and statistical analysis

Data was plotted and analysed using Graphpad version 10.4.0. Images were analysed using ImageJ software (Schneider et al., 2012). Figures were designed using Affinity Designer 2.6.2.

## Authors contributions

This work was conceptualised by M.P., who initiated the functional comparison between SymRK from *Solanum lycopersicum* and *Lotus japonicus,* as well as the domain-swap experiments. Biochemical experiments were designed by M.S. and K.P. M.S. performed nodule and primordia cross-sections, prepared samples for TEM analysis, performed complementation studies with SymRK variants expressed from different promoters, tested the interaction between SymRK ICD and SINA2, SINA4 and PUB1 using FLIM-FRET, purified proteins from *E. coli*, performed all biochemical assays, analysed the results, and prepared Fig. 1B, 1E, 3, 4, 5 and Fig. S2, S3, S5, S7, S9, S10, S13, S14. A.I.S. conducted *symrk-3* mutant complementation studies and phenotypic analysis of RNS (Fig. 1A and 1D) and AM restoration (Fig. S1) and also prepared the primordium section used in Fig. 1B. M.K.R-L. generated domain swap constructs between *Sl*SymRK and *Lj*SymRK and used them to perform hairy roots complementation of *symrk-3* (Fig. 2), and described the speckled primordia on *SlSymRK/symrk-3* (Fig. 1C). J.J. carried out all Y2H assays (Fig. 4A, B; 5, and Fig. S12, S15) using constructs prepared by M.S. Experimental design for Electron microscopy (Fig. 1E and Fig. S3) was conceptualised by A.K. and microscopy and corresponding method section was done by J.B. Mass spectrometry-based proteomic analyses (Fig. S6, S13 and S14) was performed by M.A. and C.L. on samples prepared by M.S. T.F-Ø conducted phylogenetic analysis of SINA and PUB1 homologs (Fig. S11) and sequence conservation analysis of SymRK C-terminal tail across its orthologs (Fig. S4). C.B. performed the Western Blot displayed in Fig. S10 and assisted M.S. in purifying proteins expressed in *E.coli.* A first draft of the manuscript was written by M.S, and the final version was prepared with major contributions from K.P. and M.P. The manuscript was edited by M.K.R-L, T.F-Ø., J.J., M.A., A.K., and C.L. M.P. and K.P. acquired the funding.

## Supporting information

Spezzati_Supplemental_Figures

## Acknowledgments

This project has received funding from the European Research Council (ERC) under the European Union’s Seventh Framework Programme (FP7/2007-2013) under grant agreement n° 340904 (EvolvingNodules) which supported the work of I.A.S. and M.K.R-L. This work was supported by the Deutsche Forschungsgemeinschaft (DFG) – project numbers 469056651 from 2021 until 2022 and 491090170 Transregio 356 PlantMicrobe since 2023, sub-projects B06 to M.P. and B07 to K.P. C.L. and M.A, would like to thank Franziska Hackbarth for her excellent technical assistance and maintenance of mass spectrometers. The Exploris 480 mass spectrometer was funded in part by the German Research Foundation (INST 95/1435-1 FUGG). We would like to thank Claudio Gemsa for the identification of SymRK orthologs in land plants, Vidyashree Shivakumaraswamy for preparing the PUB1 construct and Jessica Folgmann for technical support. We are grateful to Uta Paszkowski and Gabriel Ferreras Garrucho for sharing the transcriptional enhancer construct. We acknowledge use of artificial intelligence tools exclusively for manuscript editing and spelling correction. Results presented in this manuscript (Fig.1 A,D and Fig.2) were previously published in a doctoral dissertation of I.A.S.

## Data availability

Electron microscopy images, fluorescent microscopy images and raw data counts of hairy roots experiments and Y2H are deposited on GitLab and accessible on this link https://gitlab.plantmicrobe.de/maria.spezzati/Sequence_adaptation_of_SymRK_intra cellular_domain.

## References

1. Abel, N.B., Norgaard, M.M.M., Hansen, S.B., Gysel, K., Diez, I.A., Jensen, O.N., Stougaard, J., and Andersen, K.R. (2024). Phosphorylation of the alpha-I motif in SYMRK drives root nodule organogenesis. Proc Natl Acad Sci U S A 121, e2311522121.

2. Abramson, J., Adler, J., Dunger, J., Evans, R., Green, T., Pritzel, A., Ronneberger, O., Willmore, L., Ballard, A.J., Bambrick, J., Bodenstein, S.W., Evans, D.A., Hung, C.C., O’Neill, M., Reiman, D., Tunyasuvunakool, K., Wu, Z., Zemgulyte, A., Arvaniti, E., Beattie, C., Bertolli, O., Bridgland, A., Cherepanov, A., Congreve, M., Cowen-Rivers, A.I., Cowie, A., Figurnov, M., Fuchs, F.B., Gladman, H., Jain, R., Khan, Y.A., Low, C.M.R., Perlin, K., Potapenko, A., Savy, P., Singh, S., Stecula, A., Thillaisundaram, A., Tong, C., Yakneen, S., Zhong, E.D., Zielinski, M., Zidek, A., Bapst, V., Kohli, P., Jaderberg, M., Hassabis, D., and Jumper, J.M. (2024). Accurate structure prediction of biomolecular interactions with AlphaFold 3. Nature 630, 493–500.

3. Antolin-Llovera, M., Ried, M.K., and Parniske, M. (2014). Cleavage of the SYMBIOSIS RECEPTOR-LIKE KINASE ectodomain promotes complex formation with Nod factor receptor 5. Curr Biol 24, 422–427.

4. Ardourel, M., Demont, N., Debelle, F., Maillet, F., de Billy, F., Prome, J.C., Denarie, J., and Truchet, G. (1994). Rhizobium meliloti lipooligosaccharide nodulation factors: different structural requirements for bacterial entry into target root hair cells and induction of plant symbiotic developmental responses. Plant Cell 6, 1357–1374.

5. Bailey, T.L. (2021). STREME: accurate and versatile sequence motif discovery. Bioinformatics 37, 2834–2840.

6. Binder, A., Lambert, J., Morbitzer, R., Popp, C., Ott, T., Lahaye, T., and Parniske, M. (2014). A modular plasmid assembly kit for multigene expression, gene silencing and silencing rescue in plants. PLoS One 9, e88218.

7. Bonfante, P., Genre, A., Faccio, A., Martini, I., Schauser, L., Stougaard, J., Webb, J., and Parniske, M. (2000). The *Lotus japonicus LjSym4* gene is required for the successful symbiotic infection of root epidermal cells. Mol Plant Microbe Interact 13, 1109–1120.

8. Capoen, W., Goormachtig, S., De Rycke, R., Schroeyers, K., and Holsters, M. (2005). *Sr*SymRK, a plant receptor essential for symbiosome formation. Proc Natl Acad Sci U S A 102, 10369–10374.

9. Cathebras, C., Gong, X., Andrade, R.E., Vondenhoff, K., Keller, J., Delaux, P.M., Hayashi, M., Griesmann, M., and Parniske, M. (2026). A novel cis-element enabled bacterial uptake by plant cells. Nat Plants 12, 140–151.

10. Chakrabarti, D., Paul, A., Bhattacharyya, S., Das, S., Molla, F., Biswas, A., and DasGupta, M. (2024). Distinct Proline residues in hinge-regions of SYMRK generates a phosphocode for releasing Malectin-like-Domain to allow progress of rhizobia-legume symbiosis at epidermal-cortical barrier. bioRxiv, 2024.2008.2014.607956.

11. Charpentier, M., Bredemeier, R., Wanner, G., Takeda, N., Schleiff, E., and Parniske, M. (2008). *Lotus japonicus* CASTOR and POLLUX are ion channels essential for perinuclear calcium spiking in legume root endosymbiosis. Plant Cell 20, 3467–3479.

12. Chen, L., Maes, M., Cochran, A.M., Avila, J.R., Derbyshire, P., Sklenar, J., Haas, K.M., Villen, J., Menke, F.L.H., and Torii, K.U. (2025). Preventing inappropriate signals pre– and post-ligand perception by a toggle switch mechanism of ERECTA. Proc Natl Acad Sci U S A 122, e2420196122.

13. Chiasson, D., Gimenez-Oya, V., Bircheneder, M., Bachmaier, S., Studtrucker, T., Ryan, J., Sollweck, K., Leonhardt, H., Boshart, M., Dietrich, P., and Parniske, M. (2019). A unified multi-kingdom Golden Gate cloning platform. Sci Rep 9, 10131.

14. Crooks, G.E., Hon, G., Chandonia, J.M., and Brenner, S.E. (2004). WebLogo: a sequence logo generator. Genome Res 14, 1188–1190.

15. Demchenko, K., Winzer, T., Stougaard, J., Parniske, M., and Pawlowski, K. (2004). Distinct roles of *Lotus japonicus SYMRK* and *SYM15* in root colonization and arbuscule formation. New Phytol 163, 381–392.

16. Den Herder, G., Yoshida, S., Antolin-Llovera, M., Ried, M.K., and Parniske, M. (2012). *Lotus japonicus* E3 ligase SEVEN IN ABSENTIA4 destabilizes the symbiosis receptor-like kinase SYMRK and negatively regulates rhizobial infection. Plant Cell 24, 1691–1707.

17. Dixon, R.O.D. (1964). The structure of infection threads, bacteria and bacteroids in pea and clover root nodules. Archiv für Mikrobiologie 48, 166–178.

18. Doyle, J., Doyle, J., Ballenger, J., Dickson, E., Kajita, T., and Ohashi, H. (1997). A phylogeny of the chloroplast gene *rbcL* in the Leguminosae: taxonomic correlations and insights into the evolution of nodulation. Am J Bot 84, 541.

19. Doyle, J.J. (2011). Phylogenetic perspectives on the origins of nodulation. Mol Plant Microbe Interact 24, 1289–1295.

20. Emms, D.M., and Kelly, S. (2019). OrthoFinder: phylogenetic orthology inference for comparative genomics. Genome Biol 20, 238.

21. Engler, C., Kandzia, R., and Marillonnet, S. (2008). A one pot, one step, precision cloning method with high throughput capability. PLoS One 3, e3647.

22. Frank, M., Fechete, L.I., Tedeschi, F., Nadzieja, M., Norgaard, M.M.M., Montiel, J., Andersen, K.R., Schierup, M.H., Reid, D., and Andersen, S.U. (2023). Single-cell analysis identifies genes facilitating rhizobium infection in Lotus japonicus. Nat Commun 14, 7171.

23. Froger, A., and Hall, J.E. (2007). Transformation of plasmid DNA into E. coli using the heat shock method. J Vis Exp, 253.

24. Gamborg, O.L., Miller, R.A., and Ojima, K. (1968). Nutrient requirements of suspension cultures of soybean root cells. Exp Cell Res 50, 151–158.

25. Handberg, K., and Stougaard, J. (1992). *Lotus japonicus*, an autogamous, diploid legume species for classical and molecular genetics. The Plant Journal 2, 487–496.

26. Hoagland, D.R.A., D.I. (1938). The water-culture method for growing plants without soil. Circular California Agricultural Experiment Station 347.

27. Ivanov, S., and Harrison, M.J. (2024). Receptor-associated kinases control the lipid provisioning program in plant-fungal symbiosis. Science 383, 443–448.

28. Jordan, D.C., Grinyer, I., and Coulter, W.H. (1963). Electron microscopy of infection threads and bacteria in young root nodules of *Medicago sativa*. J Bacteriol 86, 125–137.

29. Katoh, K., Misawa, K., Kuma, K., and Miyata, T. (2002). MAFFT: a novel method for rapid multiple sequence alignment based on fast Fourier transform. Nucleic Acids Res 30, 3059–3066.

30. Kereszt, A., Mergaert, P., and Kondorosi, E. (2011). Bacteroid development in legume nodules: evolution of mutual benefit or of sacrificial victims? Mol Plant Microbe Interact 24, 1300–1309.

31. Keymer, A., Pimprikar, P., Wewer, V., Huber, C., Brands, M., Bucerius, S.L., Delaux, P.M., Klingl, V., Ropenack-Lahaye, E.V., Wang, T.L., Eisenreich, W., Dormann, P., Parniske, M., and Gutjahr, C. (2017). Lipid transfer from plants to arbuscular mycorrhiza fungi. Elife 6.

32. Kistner, C., and Parniske, M. (2002). Evolution of signal transduction in intracellular symbiosis. Trends Plant Sci 7, 511–518.

33. Kistner, C., Winzer, T., Pitzschke, A., Mulder, L., Sato, S., Kaneko, T., Tabata, S., Sandal, N., Stougaard, J., Webb, K.J., Szczyglowski, K., and Parniske, M. (2005). Seven *Lotus japonicus* genes required for transcriptional reprogramming of the root during fungal and bacterial symbiosis. Plant Cell 17, 2217–2229.

34. Kozlov, A.M., Darriba, D., Flouri, T., Morel, B., and Stamatakis, A. (2019). RAxML-NG: a fast, scalable and user-friendly tool for maximum likelihood phylogenetic inference. Bioinformatics 35, 4453–4455.

35. Letunic, I., and Bork, P. (2024). Interactive Tree of Life (iTOL) v6: recent updates to the phylogenetic tree display and annotation tool. Nucleic Acids Res 52, W78–W82.

36. Li, H., Chen, M., Duan, L., Zhang, T., Cao, Y., and Zhang, Z. (2018). Domain swap approach reveals the critical roles of different domains of symrk in root nodule symbiosis in *Lotus japonicus*. Front Plant Sci 9, 697.

37. Limpens, E., Mirabella, R., Fedorova, E., Franken, C., Franssen, H., Bisseling, T., and Geurts, R. (2005). Formation of organelle-like N2-fixing symbiosomes in legume root nodules is controlled by DMI2. Proc Natl Acad Sci U S A 102, 10375–10380.

38. Liu, J., Deng, J., Zhu, F., Li, Y., Lu, Z., Qin, P., Wang, T., and Dong, J. (2018). The *Mt*DMI2-*Mt*PUB2 negative feedback loop plays a role in nodulation homeostasis. Plant Physiol 176, 3003–3026.

39. Lu, D., Lin, W., Gao, X., Wu, S., Cheng, C., Avila, J., Heese, A., Devarenne, T.P., He, P., and Shan, L. (2011). Direct ubiquitination of pattern recognition receptor FLS2 attenuates plant innate immunity. Science 332, 1439–1442.

40. Luginbuehl, L.H., Menard, G.N., Kurup, S., Van Erp, H., Radhakrishnan, G.V., Breakspear, A., Oldroyd, G.E.D., and Eastmond, P.J. (2017). Fatty acids in arbuscular mycorrhizal fungi are synthesized by the host plant. Science 356, 1175–1178.

41. MacLean, B., Tomazela, D.M., Shulman, N., Chambers, M., Finney, G.L., Frewen, B., Kern, R., Tabb, D.L., Liebler, D.C., and MacCoss, M.J. (2010). Skyline: an open source document editor for creating and analyzing targeted proteomics experiments. Bioinformatics 26, 966–968.

42. Markmann, K., Giczey, G., and Parniske, M. (2008). Functional adaptation of a plant receptor-kinase paved the way for the evolution of intracellular root symbioses with bacteria. PLoS Biol 6, e68.

43. Mbengue, M., Camut, S., de Carvalho-Niebel, F., Deslandes, L., Froidure, S., Klaus-Heisen, D., Moreau, S., Rivas, S., Timmers, T., Herve, C., Cullimore, J., and Lefebvre, B. (2010). The *Medicago truncatula* E3 ubiquitin ligase PUB1 interacts with the LYK3 symbiotic receptor and negatively regulates infection and nodulation. Plant Cell 22, 3474–3488.

44. Meng, E.C., Goddard, T.D., Pettersen, E.F., Couch, G.S., Pearson, Z.J., Morris, J.H., and Ferrin, T.E. (2023). UCSF ChimeraX: Tools for structure building and analysis. Protein Sci 32, e4792.

45. Ngou, B.P.M., Wyler, M., Schmid, M.W., Kadota, Y., and Shirasu, K. (2024). Evolutionary trajectory of pattern recognition receptors in plants. Nat Commun 15, 308.

46. Offringa, I.A., Melchers, L.S., Regensburg-Tuink, A.J., Costantino, P., Schilperoort, R.A., and Hooykaas, P.J. (1986). Complementation of Agrobacterium tumefaciens tumor-inducing aux mutants by genes from the T(R)-region of the Ri plasmid of Agrobacterium rhizogenes. Proc Natl Acad Sci U S A 83, 6935–6939.

47. Perry, J.A., Wang, T.L., Welham, T.J., Gardner, S., Pike, J.M., Yoshida, S., and Parniske, M. (2003). A TILLING reverse genetics tool and a web-accessible collection of mutants of the legume Lotus japonicus. Plant Physiol 131, 866–871.

48. Ried, M.K., Antolin-Llovera, M., and Parniske, M. (2014). Spontaneous symbiotic reprogramming of plant roots triggered by receptor-like kinases. Elife 3.

49. Rodriguez-Furlan, C., Minina, E.A., and Hicks, G.R. (2019). Remove, Recycle, Degrade: regulating plasma membrane protein accumulation. Plant Cell 31, 2833–2854.

50. Saha, S., Dutta, A., Bhattacharya, A., and DasGupta, M. (2014). Intracellular catalytic domain of symbiosis receptor kinase hyperactivates spontaneous nodulation in absence of rhizobia. Plant Physiol 166, 1699–1708.

51. Saha, S., Paul, A., Herring, L., Dutta, A., Bhattacharya, A., Samaddar, S., Goshe, M.B., and DasGupta, M. (2016). Gatekeeper tyrosine phosphorylation of SYMRK is essential for synchronizing the epidermal and cortical responses in root nodule symbiosis. Plant Physiol 171, 71–81.

52. Schauser, L., Roussis, A., Stiller, J., and Stougaard, J. (1999). A plant regulator controlling development of symbiotic root nodules. Nature 402, 191–195.

53. Schneider, C.A., Rasband, W.S., and Eliceiri, K.W. (2012). NIH Image to ImageJ: 25 years of image analysis. Nat Methods 9, 671–675.

54. Schüβler, A., Schwarzott, D., and Walker, C. (2001). A new fungal phylum, the Glomeromycota: phylogeny and evolution* *Dedicated to Manfred Kluge (Technische Universität Darmstadt) on the occasion of his retirement. Mycological Research 105, 1413–1421.

55. Sevin-Pujol, A., Sicard, M., Rosenberg, C., Auriac, M.C., Lepage, A., Niebel, A., Gough, C., and Bensmihen, S. (2017). Development of a GAL4-VP16/UAS trans-activation system for tissue specific expression in *Medicago truncatula*. PLoS One 12, e0188923.

56. Shevchenko, A., Tomas, H., Havlis, J., Olsen, J.V., and Mann, M. (2006). In-gel digestion for mass spectrometric characterization of proteins and proteomes. Nat Protoc 1, 2856–2860.

57. Singh, S., Katzer, K., Lambert, J., Cerri, M., and Parniske, M. (2014). CYCLOPS, a DNA-binding transcriptional activator, orchestrates symbiotic root nodule development. Cell Host Microbe 15, 139–152.

58. Stegmann, M., Anderson, R.G., Ichimura, K., Pecenkova, T., Reuter, P., Zarsky, V., McDowell, J.M., Shirasu, K., and Trujillo, M. (2012). The ubiquitin ligase PUB22 targets a subunit of the exocyst complex required for PAMP-triggered responses in Arabidopsis. Plant Cell 24, 4703–4716.

59. Stracke, S., Kistner, C., Yoshida, S., Mulder, L., Sato, S., Kaneko, T., Tabata, S., Sandal, N., Stougaard, J., Szczyglowski, K., and Parniske, M. (2002). A plant receptor-like kinase required for both bacterial and fungal symbiosis. Nature 417, 959–962.

60. Tyanova, S., Temu, T., and Cox, J. (2016). The MaxQuant computational platform for mass spectrometry-based shotgun proteomics. Nat Protoc 11, 2301–2319.

61. Vernié, T., Camut, S., Camps, C., Rembliere, C., de Carvalho-Niebel, F., Mbengue, M., Timmers, T., Gasciolli, V., Thompson, R., le Signor, C., Lefebvre, B., Cullimore, J., and Herve, C. (2016). PUB1 interacts with the receptor kinase DMI2 and negatively regulates rhizobial and arbuscular mycorrhizal symbioses through its ubiquitination activity in *Medicago truncatula*. Plant Physiol 170, 2312–2324.

62. Vierheilig, H., Coughlan, A.P., Wyss, U., and Piche, Y. (1998). Ink and vinegar, a simple staining technique for arbuscular-mycorrhizal fungi. Appl Environ Microbiol 64, 5004–5007.

63. Wang, Z.Y., Seto, H., Fujioka, S., Yoshida, S., and Chory, J. (2001). BRI1 is a critical component of a plasma-membrane receptor for plant steroids. Nature 410, 380–383.

64. Wei, Q., Qiang, F., Zhang, X., Wang, F., Liu, J., Fan, H., Liu, M., Li, G., and Wu, G. (2025). Identification of inhibitions of the C-terminal tails of canonical (but not orphan) brassinosteroid receptor kinases reveals intramolecular antagonistic coevolution. Int J Biol Macromol 332, 148638.

65. Yoshida, S., and Parniske, M. (2005). Regulation of plant symbiosis receptor kinase through serine and threonine phosphorylation. J Biol Chem 280, 9203–9209.

