## Supplementary material for "Sequence adaptations in the intracellular domain of Symbiosis receptor-like kinase (SymRK) promoted infection thread progression in root nodule primordia": Spezzati_Supplemental_Figures

### Supplementary Figures

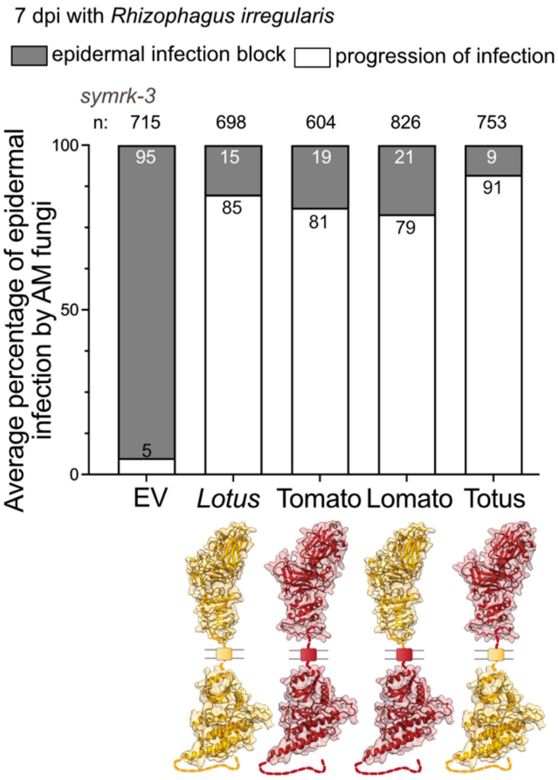


**Supplementary Figure 1. *Sl*SymRK and its chimeric variants restore progression of AM fungal infection across the root epidermis in the *Lotus symrk-3* mutant.**

The average percentage of early infection structures was quantified on *symrk-3* roots expressing the corresponding SymRK chimeras (depicted below the plot) seven days post-inoculation (dpi) with *Rhizophagus irregularis*. All tested SymRK variants complemented *symrk-3* for the progression of epidermal infection to sub-epidermal and cortical cells. The empty vector (EV) was used as negative control with 95 % aborted epidermal entries. Statistical significance was evaluated by two-way ANOVA followed by Šídák 's multiple comparison test. Note that there was no statistically significant difference in the percentage of epidermal infection events among the different variants of *Sl*SymRK compared to *Lotus* SymRK. For each genetic construct, 13 to 15 plants were analysed. n: number of counted infection structures. The depicted 3D structures of SymRK *Lotus* and tomato were predicted using AlphaFold 3.

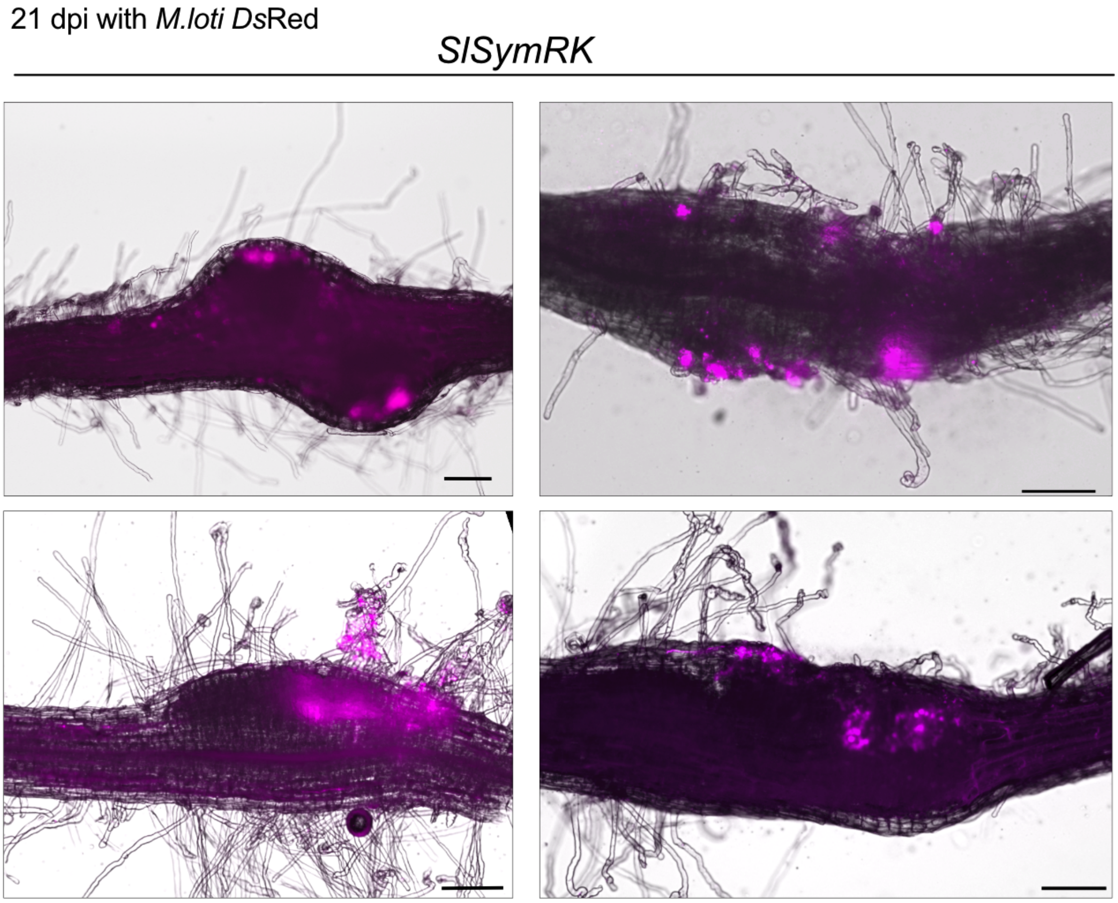


**Supplementary Figure 2. Speckled primordia on *L. japonicus symrk-3* hairy roots transformed with *proLjSymRK:SlSymRK***

Merged images (bright field and *Ds*Red signal (magenta)) of speckled primordia on transformed *symrk-3* mutant roots transformed with *Sl*SymRK driven by *Lotus* SymRK promoter 21 days post inoculation with *M. loti Ds*Red. Note that rhizobia accumulate in the outer cell layers of the speckled primordia (magenta) but not the central tissue. Scale bar: 150 µm.

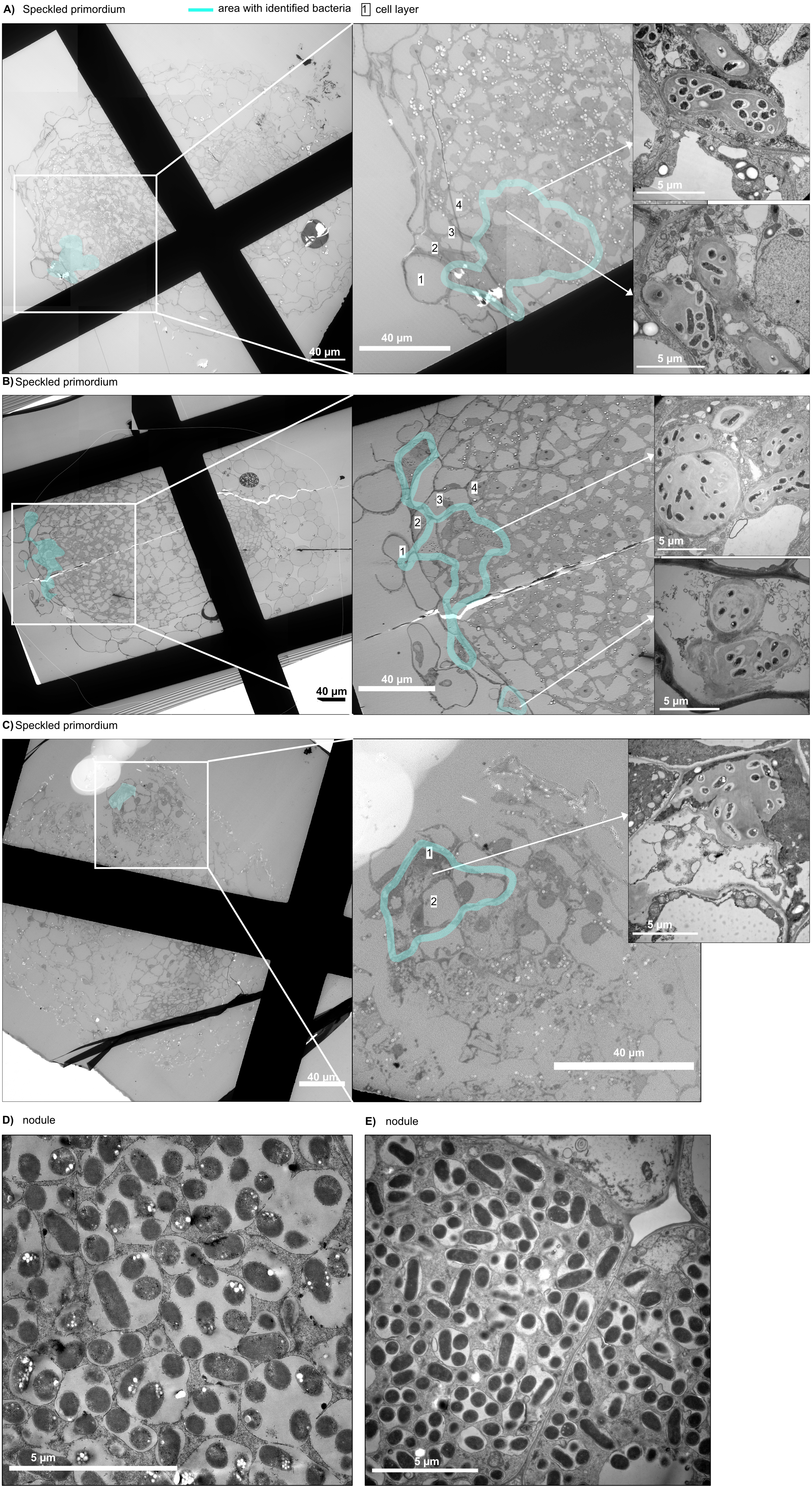
A) Speckled primordium
area with identified bacteria
1 cell layer
40 µm
4
3
2
1
40 µm
5 µm
5 µm
B) Speckled primordium
40 µm
4
3
2
1
40 µm
5 µm
5 µm
C) Speckled primordium
40 µm
1
2
40 µm
5 µm
D) nodule
5 µm
E) nodule
5 µm

**Supplementary Figure 3. Infection threads in speckled primordia on *SlSymRK/symrk-3*.**

A) - C) Electron micrographs of three distinct speckled primordia cross sections observed on *proLjSymRK:SlSymRK/symrk-3* roots 21 dpi with *M.loti Ds*Red. On the left, overview image of a section on the sample support grid of the electron microscope. The cells in which bacteria were observed are highlighted in cyan. In succession to the right, increased magnification of area of plant cells containing bacteria and two representative cells that contained bacteria. Note that the bacteria are surrounded by a structure that resembles infection threads and that bacteria were observed in this case in the first four outer cell layers but not in the central tissue of the primordium. The numbers in the white square indicate consecutive cell layers. Scale bar: 40 μm (left), 5 μm (right).

D) and E) electron micrographs of two nodule samples cross sections observed on *Lj*SymRK/*symrk-3* roots 21 dpi with *M.loti Ds*Red. Images are showing the central tissue of the nodule where symbiosomes can be observed. Scale bar: 5 μm.

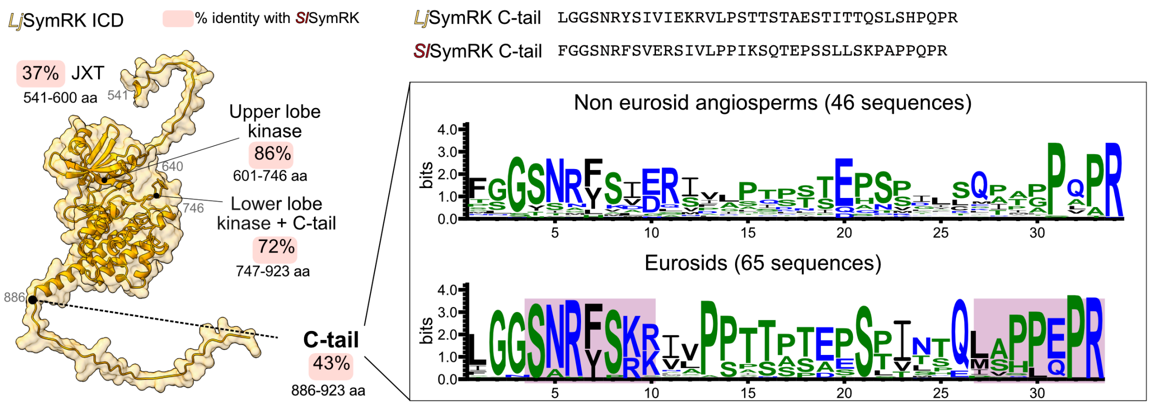


**Supplementary Figure 4. The unstructured C-terminal portion of SymRK is variable among its orthologs.**

Left: AlphaFold 3 model of *Lj*SymRK ICD with annotated percent amino acid sequence identity between *Lotus* and tomato SymRK calculated for four regions of their ICDs. The borders of the subdomains of *Lj*SymRK ICD are indicated by a black dot. Note the low percentage identity for the unstructured juxtamembrane (JXT) and C-terminal (C-tail) regions. Right: Amino acid sequence of the C-tail from *Lj*SymRK and *Sl*SymRK. Below: Amino acid sequence logos obtained from the alignment of C-tail regions of SymRK orthologs from Eurosids species and non-Eurosids angiosperms species. Motifs that are enriched in the Eurosids clade relative to other angiosperms are highlighted in red boxes. Sequence logos were generated using WebLogo (Crooks et al., 2004).

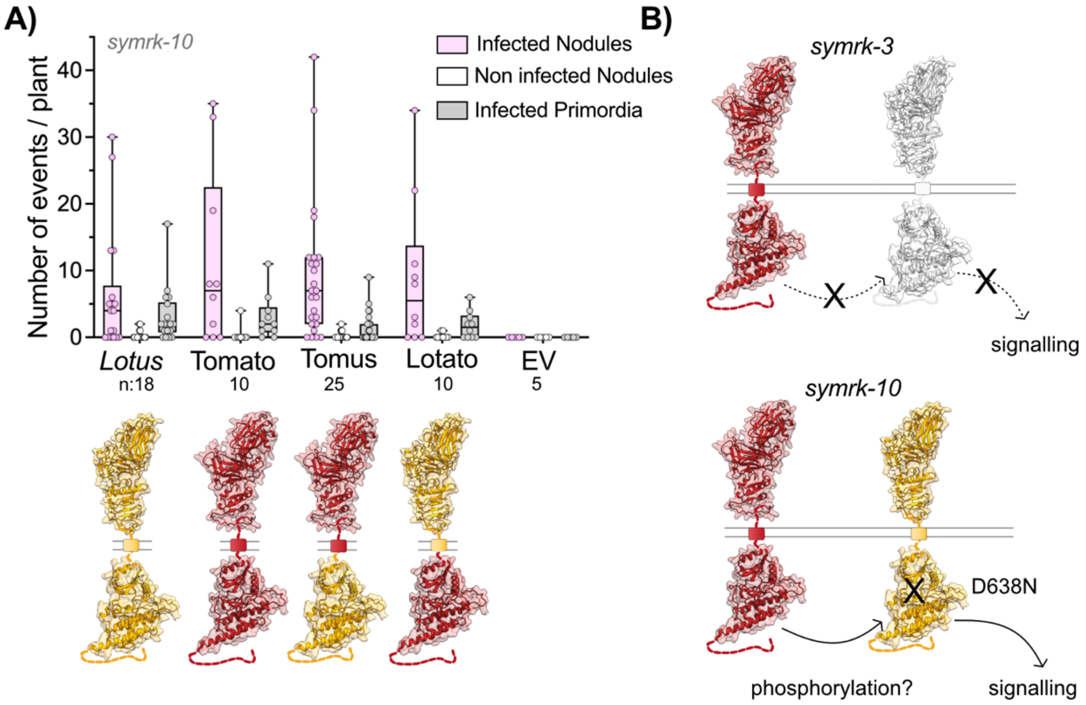


**Supplementary Figure 5. *Sl*SymRK restores RNS in the *symrk-10* mutant.**

A) Different variants of *SymRK* (*Lotus*, Tomato, Tomus and Lotato) expressed under the native *SymRK* promoter*,* complemented the kinase-dead mutant *symrk-10*. Notably, no significant differences were observed for number of nodules and primordia for Tomato, Tomus, Lotato compared to *Lotus* SymRK, in contrast to Empty Vector (EV) negative control. Statistical significance was assessed by two-way ANOVA followed by Dunnet's multiple comparison test for the different categories. n: number of transformed plants analysed. All phenotypes were scored at 21 dpi with *M. loti Ds*Red.

B) Model illustrating the hypothesized molecular events in the *symrk-3* (above) and *symrk-10* (below) mutant complemented with *Sl*SymRK. The *symrk-10* allele encodes a *Lj*SymRK with a an amino-acid replacement in the DFG motif (D638N). Note the difference between *symrk-3* (no *Lj*SymRK present) and *symrk-10* (*Lj*SymRK present but catalytically inactive).

The 3D structures of SymRK *Lotus* and Tomato were predicted using AlphaFold 3.

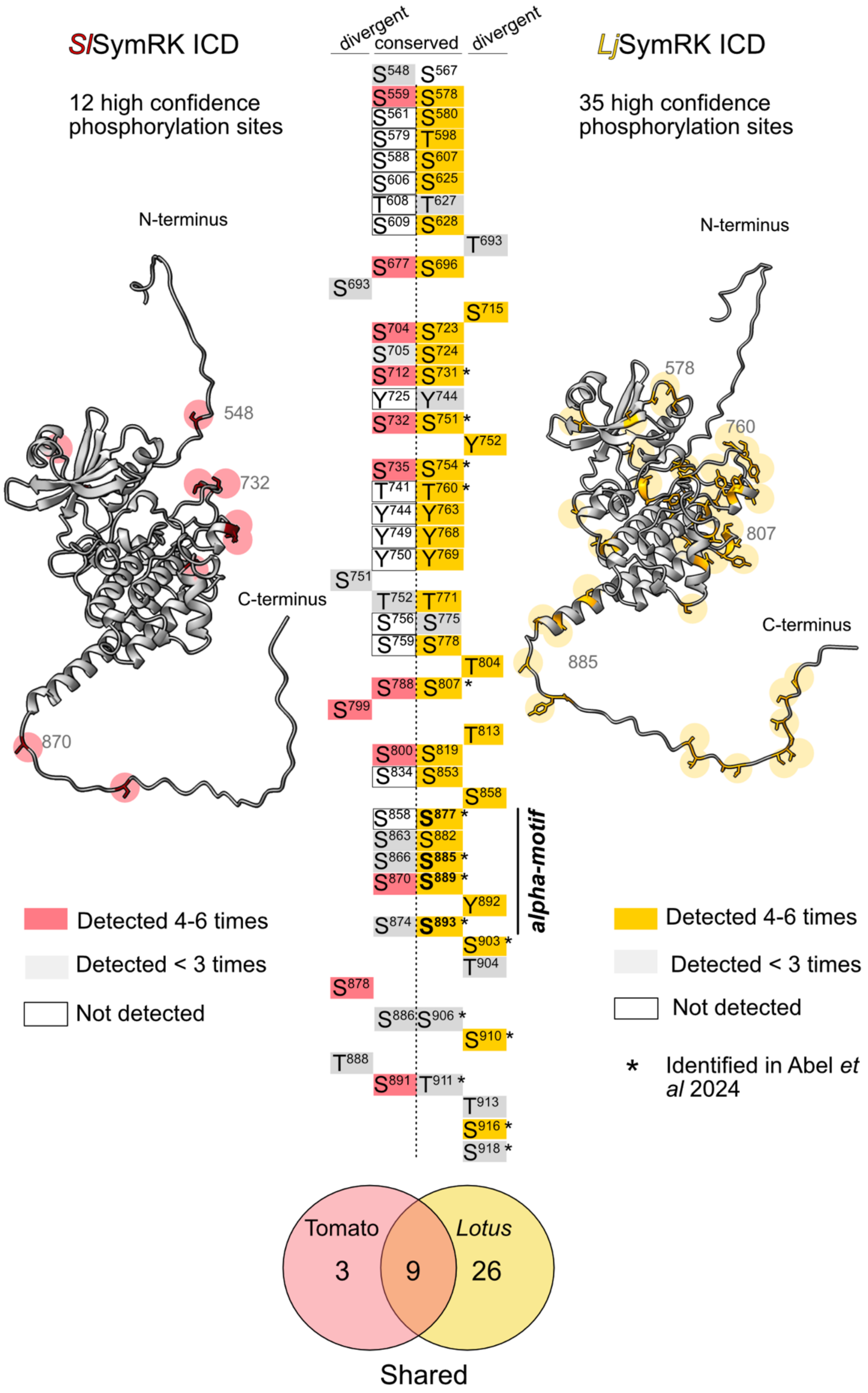

SlSymRK ICD
12 high confidence phosphorylation sites
LjSymRK ICD
35 high confidence phosphorylation sites
divergent
conserved
divergent
N-terminus
C-terminus
548
732
870
578
760
807
885
S548 S567
S559 S578
S561 S580
S579 T598
S588 S607
S606 S625
T608 T627
S609 S628
T693
S677 S696
S693
S715
S704 S723
S705 S724
S712 S731 *
Y725 Y744
S732 S751 *
Y752
S735 S754 *
T741 T760 *
Y744 Y763
Y749 Y768
Y750 Y769
S751
T752 T771
S756 S775
S759 S778
T804
S788 S807 *
S799
T813
S800 S819
S834 S853
S858
S858 S877 *
S863 S882
S866 S885 *
S870 S889 *
Y892
S874 S893 *
S903 *
T904
alpha-motif
S878
S886 S906 *
S910 *
T888
S891 T911 *
T913
S916 *
S918 *
Detected 4-6 times
Detected < 3 times
Not detected
Detected 4-6 times
Detected < 3 times
Not detected
* Identified in Abel et al 2024
Tomato
3
9
Lotus
26
Shared

**Supplementary Figure 6. The intracellular domains of *Lotus* and Tomato SymRK are auto-phosphorylated *in vitro.***

The ICDs of tomato (red) and *Lotus* (yellow) SymRK were expressed and purified from *E. coli,* incubated in a buffer containing ATP and $MgCl_2$ to enable for kinase activity (see Materials and Methods) and subjected to liquid-chromatography tandem mass spectrometry (LC-MS/MS) analysis to detect protein phosphorylation. Phosphorylation sites were filtered for sites that were reproducibly detectable in at least four out of six technical replicates. Based on this threshold, 12 high-confidence auto-phosphorylation sites were identified on tomato SymRK (shown in red), while thirty-five were found on *Lotus* SymRK (highlighted in yellow). Sites that were found in less than three replicates are shown in grey, and sites that were conserved but not detected are shown in white boxes. The identified sites were confirmed to be surface-exposed and mapped onto the corresponding AlphaFold 3-predicted structural model of SymRK ICDs from tomato (left) and *Lotus* (right). In both structural models, side chains of modified amino acids are shown, and highlighted in red for tomato and yellow for *Lotus*. Residue positions of phosphorylated Serine (S), Threonine (T) and Tyrosine (Y) that are conserved in both SymRK variants based on amino acid sequence alignment are listed in the middle, while divergent phosphorylated residues are listed at the left and right of the centre. Venn diagram: overlap between phosphorylation sites detected on tomato and *Lotus* SymRK ICD (considering only high-confidence sites). Phosphorylation sites within the alpha-motif of *Lotus* SymRK that are functionally important in RNS are highlighted in bold in the Figure (Abel et al., 2024). Asterisks correspond to phosphorylation sites also identified by Abel et al., 2024. Sites S578, 580, 751, 754 and 889 were detected with low confidence (not in all replicates) on the kinase dead *Lj*SymRK$^{K662E}$ (Supp. Table 3).

**A)**

| Promoter name | Description |
|---|---|
| *proSymRK* | endogenous SymRK promoter 4.9 Kb |
| *proSym$^{enhanced}$* | SymRK promoter + VP16-Gal4-5xUAS_pro35min |
| *proUbi* | *L. japonicus* Ubiquitin promoter |

**B)**

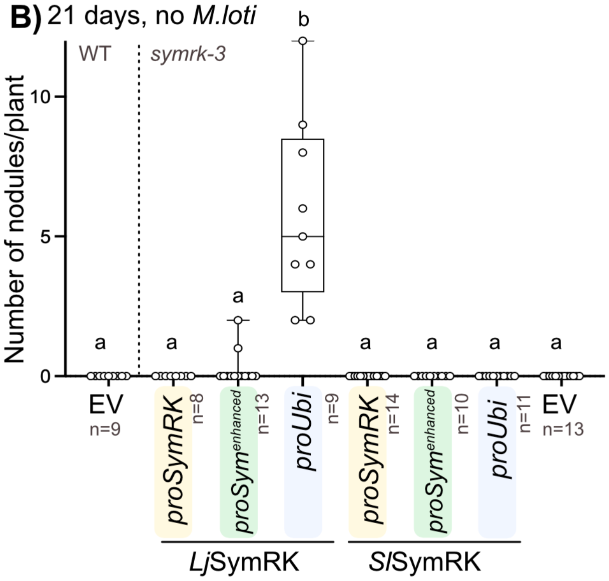


**Supplementary Figure 7. *Lotus* SymRK does, but Tomato SymRK does not induce spontaneous root nodules when expressed under a ubiquitin promoter.**

A) Promoters used for driving *Lotus* and tomato *SymRK* expression.

B) Number of spontaneous nodules observed on roots of *symrk-3* expressing the *Lotus* or tomato orthologs of *SymRK* under one of the promoters described in A). EV in *L. japonicus* Gifu (WT) was used as positive control. Spontaneous nodules were quantified 21 days after growing plants in Weck Jars in the absence of *M. loti*. Statistical significance was assessed using One-way ANOVA followed by Tukey's multiple comparisons test. Significance between samples (p-value < 0.05) is indicated by different letters. n number of plants.

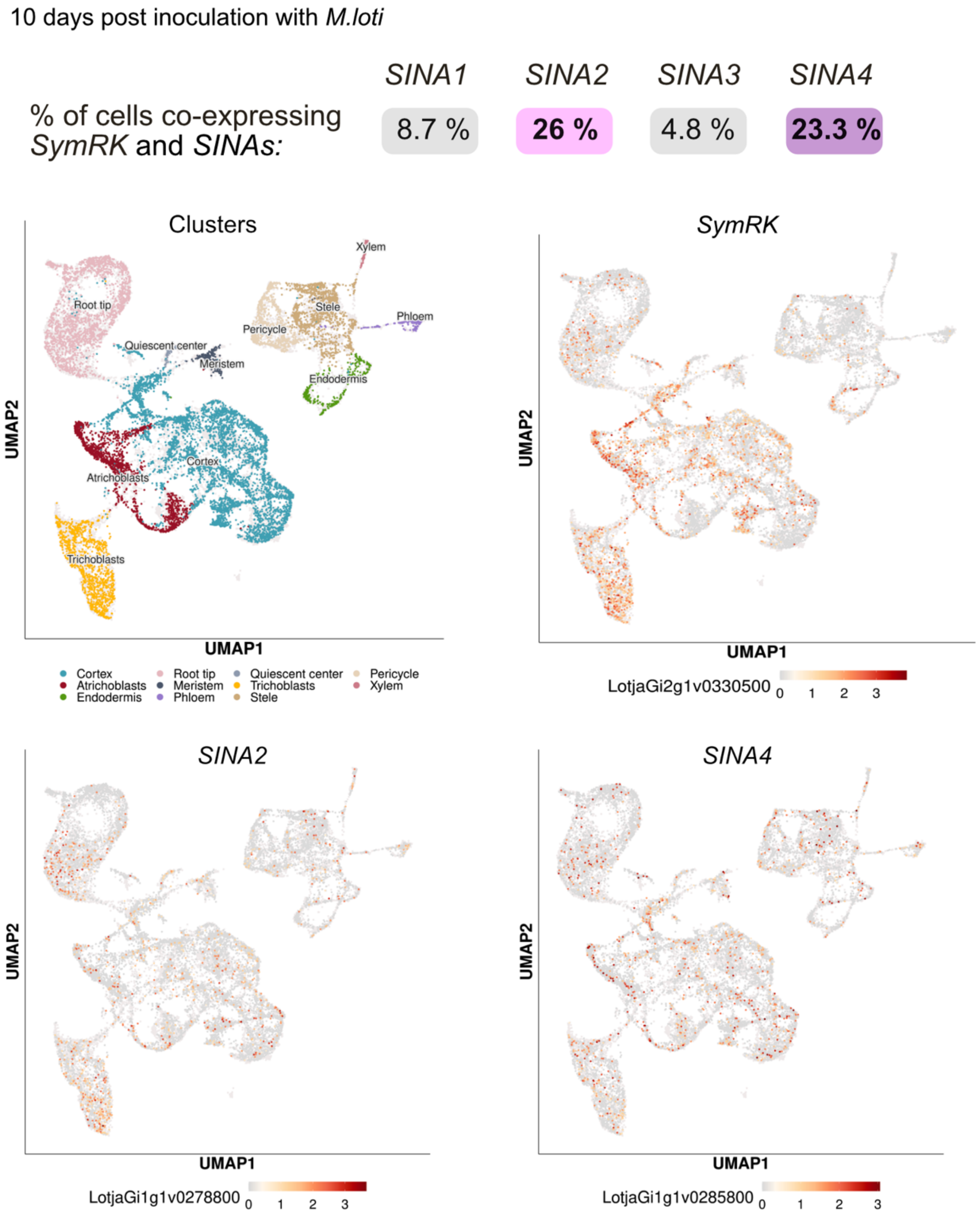


**Supplementary Figure 8. Single-cell expression pattern of *SINAs* and *SymRK* in *Lotus japonicus* roots inoculated with *M. loti*.**

UMAP plots represent coloured root clusters (left panel) and expression of candidate genes (*SymRK, SINA2* and *SINA4*). Percentage of cells in which transcripts of both *SymRK* and *SINA* members have been detected are displayed above the UMAP plots. Data derived from Frank *et al*., 2023.

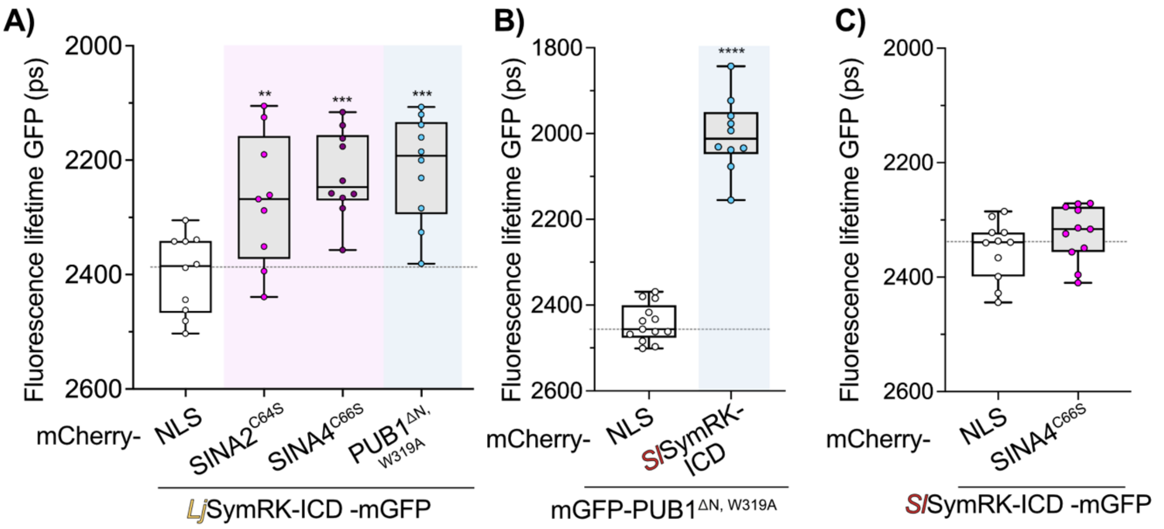


**Supplementary Figure 9. Differential interaction between SymRK and SINAs detected by FLIM-FRET *in vivo*.**

A) FLIM-FRET proximity assays conducted with *Lotus* SymRK and SINA2/4 and PUB1 E3 ligases. Fluorescence lifetime of mGFP was measured in *Nicotiana benthamiana* nuclei co-expressing *Lj*SymRK-ICD-mGFP as a donor and SINA2/4 or PUB1 fused to mCherry as an acceptor protein. E3 ligases were expressed in their inactive forms by introducing point mutations in enzymatically active sites: SINA2$^{C64S}$, SINA4$^{C66S}$ and PUB1$^{\Delta N,W319A}$. All proteins were expressed under the CaMV *35S* promoter and targeted to the nucleus by fusing a nuclear localisation signal (NLS) to their N-terminus (SymRK-ICD) or C-terminus (SINA2/4 and PUB1). NLS-mCherry was used as a negative control. Note that *Lj*SymRK-ICD interacts with all tested E3 ligases as indicated by the reduced lifetime of mGFP fluorescence. Statistical significance was assessed by one-way ANOVA followed by Dunnett's multiple comparison test; *** - p = 0.002 - 0.0002, ** - p = 0.03 - 0.0021.

B and C) FLIM-FRET interaction studies were conducted between tomato SymRK and PUB1$^{\Delta N,\ W319A}$ (B) or SINA4 $^{C66S}$ (C) E3 ligases. In panel B) mGFP- PUB1$^{\Delta N,W319A}$ was used as the donor and *Sl*SymRK-mCherry as the acceptor. Statistical analysis was performed using an unpaired t-test and statistical significance is indicated by **** (p-value < 0.0001). In panel C) *Sl*SymRK-ICD-mGFP was used as the donor and mCherry-SINA4$^{C66S}$ as the acceptor. Data were analysed with an unpaired t-test. No significant reduction in fluorescence lifetime was observed between *Sl*SymRK-mGFP and mCherry-SINA4 $^{C66S}$.

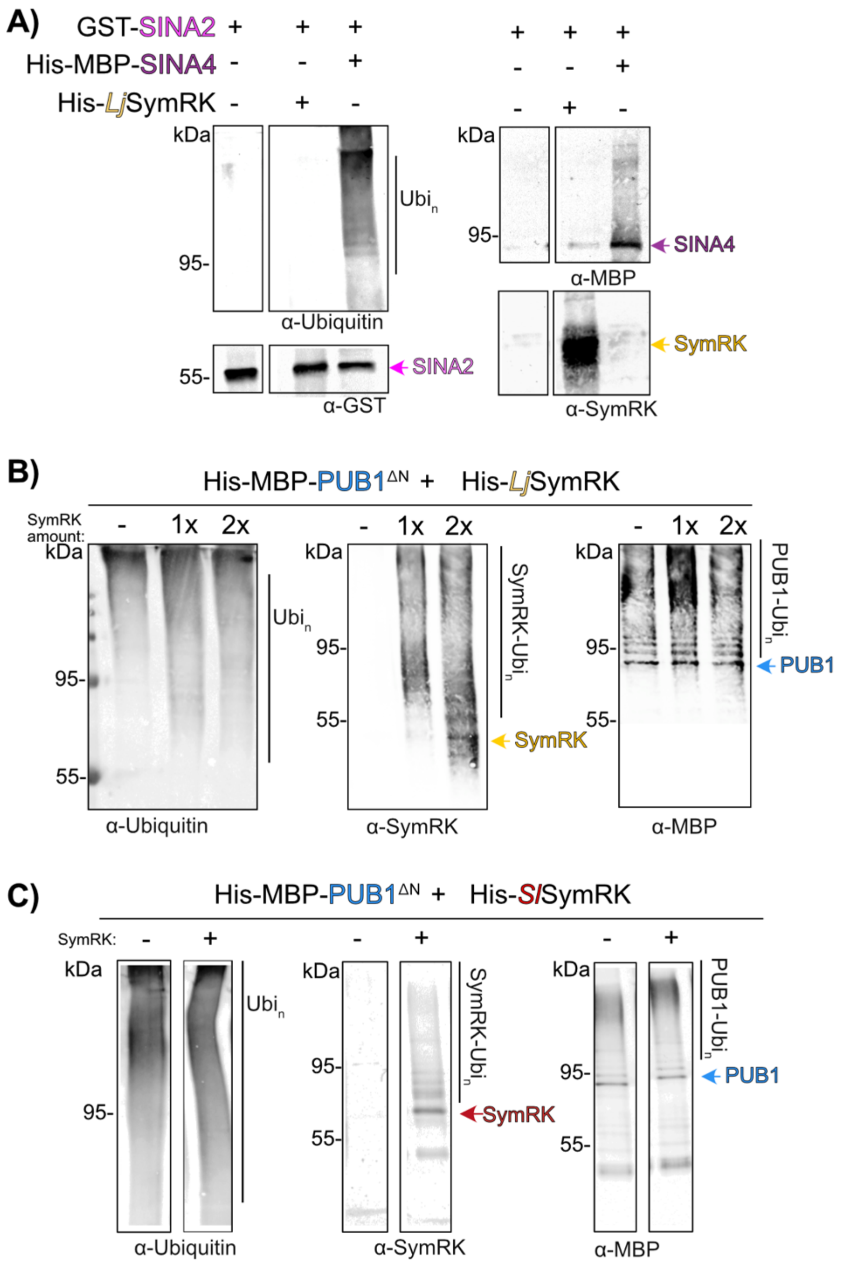


**Supplementary Figure 10. *In vitro* ubiquitinylation assays of SINA2 and PUB1$^{\Delta N}$ with *Lotus* or tomato SymRK ICD.**

A) *In vitro* ubiquitinylation assay with GST-SINA2, His-MBP-SINA4 and His-*Lj*SymRK ICD proteins purified from *E. coli*. The ubiquitinylation patterns of the proteins were analysed through Western Blot with anti-Ubiquitin, anti-SymRK, anti-GST and anti-MBP antibodies. In contrast to SINA4, GST-SINA2 did not reveal any ubiquitinylation activity *in vitro*. The yellow arrowhead marks expected size for non-modified SymRK ICD (43 kDa), while the pink and purple arrowheads mark expected size for non-modified SINA2 (61 kDa) and SINA4 (80 kDa), respectively.

B) and C) *In vitro* ubiquitinylation assay with His-MBP-PUB1$^{\Delta N}$, His-*Lj*SymRK ICD and His-*Sl*SymRK ICD proteins purified from *E. coli*. The ubiquitinylation patterns of the proteins were analysed through Western Blot with anti-Ubiquitin, anti-SymRK and anti-MBP antibodies. In B) increasing amounts of SymRK were used, as indicated above the membranes with an “x”. Note that presence of high-molecular weight signal for membranes incubated with anti-SymRK antibodies in B and C indicating *Lotus* and tomato SymRK ubiquitylation by PUB1. The yellow or red arrowhead marks expected size for non-modified SymRK ICD (43 kDa), while the blue arrowhead marks PUB1$^{\Delta N}$ (87 kDa). GST: Glutathione S-transferase, MBP: Maltose-binding Protein.

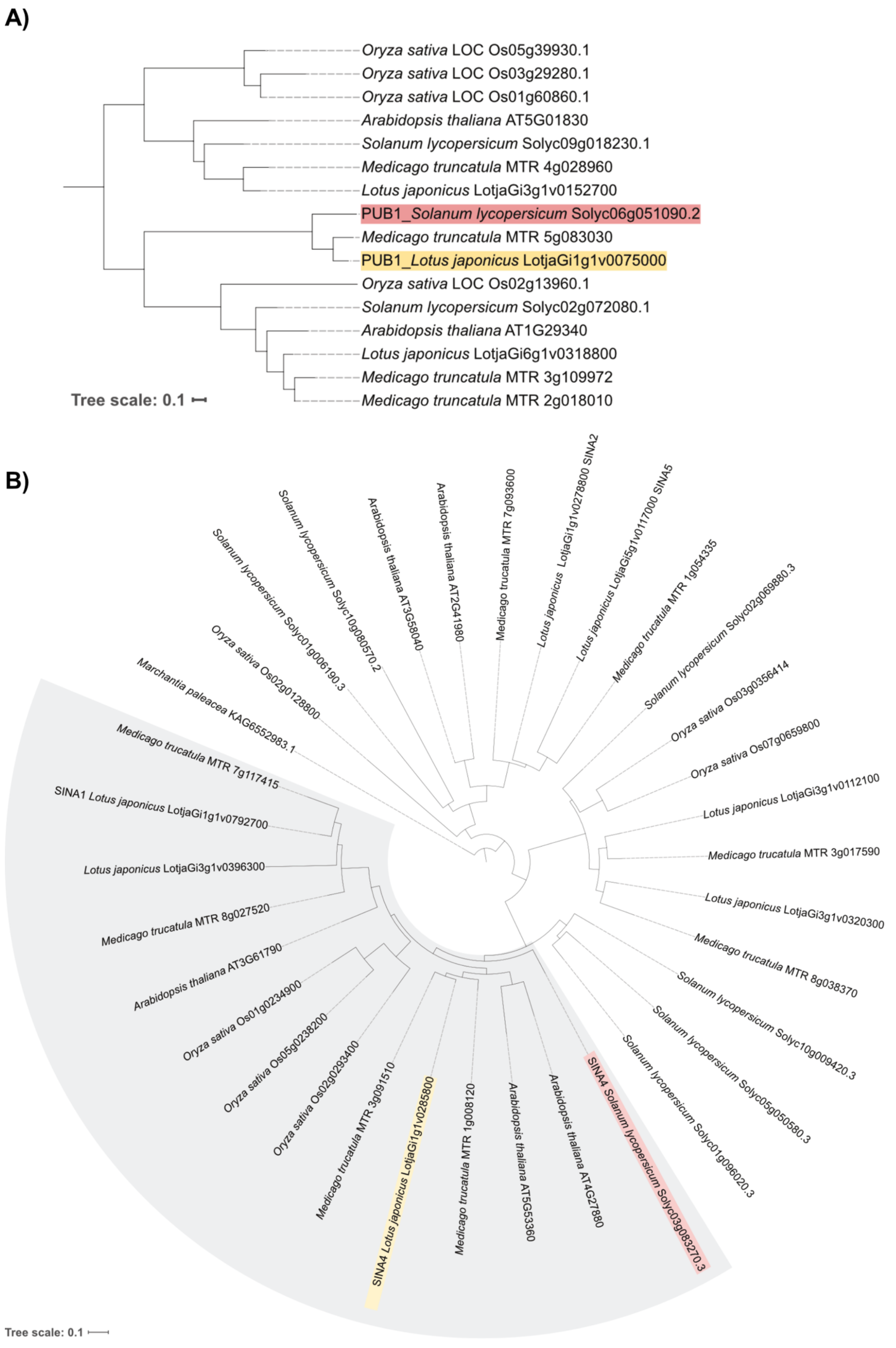
A)
Oryza sativa LOC Os05g39930.1
Oryza sativa LOC Os03g29280.1
Oryza sativa LOC Os01g60860.1
Arabidopsis thaliana AT5G01830
Solanum lycopersicum Solyc09g018230.1
Medicago truncatula MTR 4g028960
Lotus japonicus LotjaGi3g1v0152700
PUB1_Solanum lycopersicum Solyc06g051090.2
Medicago truncatula MTR 5g083030
PUB1_Lotus japonicus LotjaGi1g1v0075000
Oryza sativa LOC Os02g13960.1
Solanum lycopersicum Solyc02g072080.1
Arabidopsis thaliana AT1G29340
Lotus japonicus LotjaGi6g1v0318800
Medicago truncatula MTR 3g109972
Medicago truncatula MTR 2g018010
Tree scale: 0.1
B)
Solanum lycopersicum Solyc01g006190.3
Solanum lycopersicum Solyc10g080570.2
Arabidopsis thaliana AT3G58040
Arabidopsis thaliana AT2G41980
Medicago trucatula MTR 7g093600
Lotus japonicus LotjaGi1g1v0278800 SINA2
Lotus japonicus LotjaGi5g1v0117000 SINA5
Medicago trucatula MTR 1g054335
Solanum lycopersicum Solyc02g069880.3
Oryza sativa Os03g0356414
Oryza sativa Os07g0659800
Lotus japonicus LotjaGi3g1v0112100
Medicago trucatula MTR 3g017590
Lotus japonicus LotjaGi3g1v0320300
Medicago trucatula MTR 8g038370
Solanum lycopersicum Solyc10g009420.3
Solanum lycopersicum Solyc05g050580.3
Solanum lycopersicum Solyc01g096020.3
SINA4 Solanum lycopersicum Solyc03g083270.3
Arabidopsis thaliana AT4G27880
Arabidopsis thaliana AT5G53360
Medicago trucatula MTR 1g008120
SINA4 Lotus japonicus LotjaGi1g1v0285800
Medicago trucatula MTR 3g091510
Oryza sativa Os02g0293400
Oryza sativa Os05g0238200
Oryza sativa Os01g0234900
Arabidopsis thaliana AT3G61790
Medicago trucatula MTR 8g027520
Lotus japonicus LotjaGi3g1v0396300
SINA1 Lotus japonicus LotjaGi1g1v0792700
Medicago trucatula MTR 7g117415
Marchantia paleacea KAG6552983.1
Oryza sativa Os02g0128800
Tree scale: 0.1

**Supplementary Figure 11. Phylogenetic analysis of SINA and PUB1 homologs.**

A) Midpoint-rooted phylogenetic tree of members of the PUB1 orthogroup, filtered to include orthologs from *Lotus japonicus*, *Solanum lycopersicum*, *Medicago truncatula*, *Arabidopsis thaliana* and *Marchantia paleacea*. Tomato and *Lotus* orthologs are highlighted in red and yellow, respectively. The tree was built using RaXML-NG.

B) Tree of members of the SINA4 orthogroup, filtered to include orthologs from *Lotus japonicus*, *Solanum lycopersicum*, *Medicago truncatula*, *Arabidopsis thaliana* and *Marchantia paleacea*. The tree was built using RaXML-NG. The putative *Marchantia paleacea* ortholog was used to root the tree. Tomato and *Lotus* orthologs are highlighted in red and yellow, respectively. The clade containing *Sl*SINA4 and *Lj*SINA4, as well as *Lj*SINA1, is highlighted in grey.

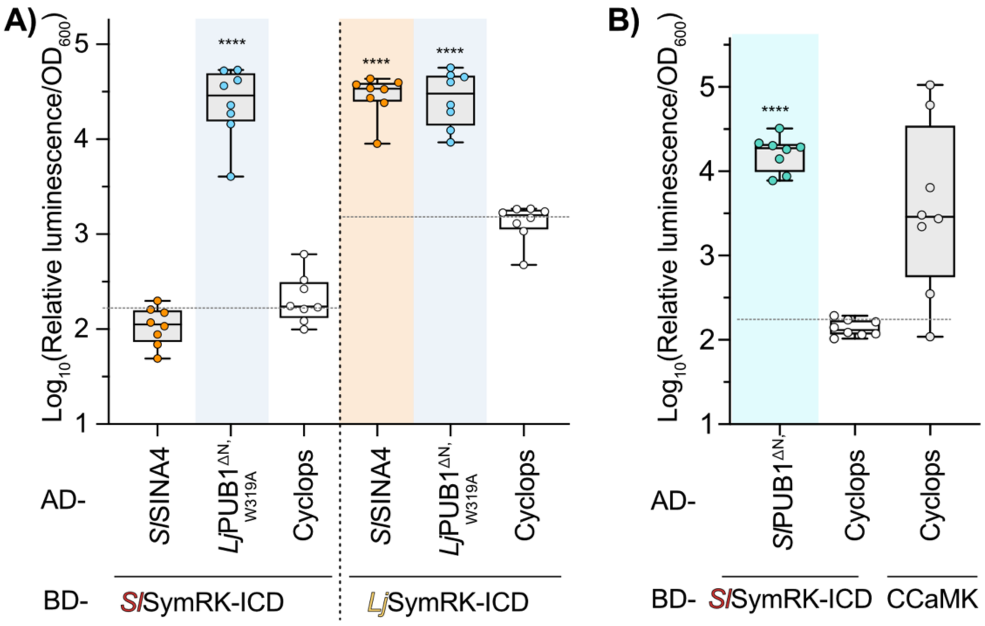


**Supplementary Figure 12. The interaction with SINA4 was detected with *Lotus* but not tomato SymRK ICD.**

A) Yeast two-hybrid interaction analysis between *Lotus* (*Lj*SymRK) or tomato ICD (*Sl*SymRK) fused to the Binding Domain (BD) and tomato SINA4 (*Sl*SINA4) or *Lj*PUB1$^{\Delta N, W319A}$ fused to the Activation Domain (AD). Cyclops-AD was used as a negative control. Note the significant signal intensity for tomato SINA4 with *Lotus* SymRK ICD, suggesting interaction, but not with tomato SymRK.

B) Tomato ortholog of *Lj*PUB1 (*Sl*PUB1$^{\Delta N}$) interacts with tomato SymRK in Yeast two-hybrid assay. Cyclops-AD was used as a negative control, while Cyclops-CCaMK was used as positive control.

In A) and B) the boxplots represent normalized luciferase signals measured for independent yeast colonies co-transformed with the indicated bait and prey constructs. A yeast strain carrying an integrated *proGal2:Luciferase* reporter was used in the studies. Statistical significance was assessed using one-way ANOVA followed by Tukey's multiple comparisons test; **** : p-value $< 0.05$.

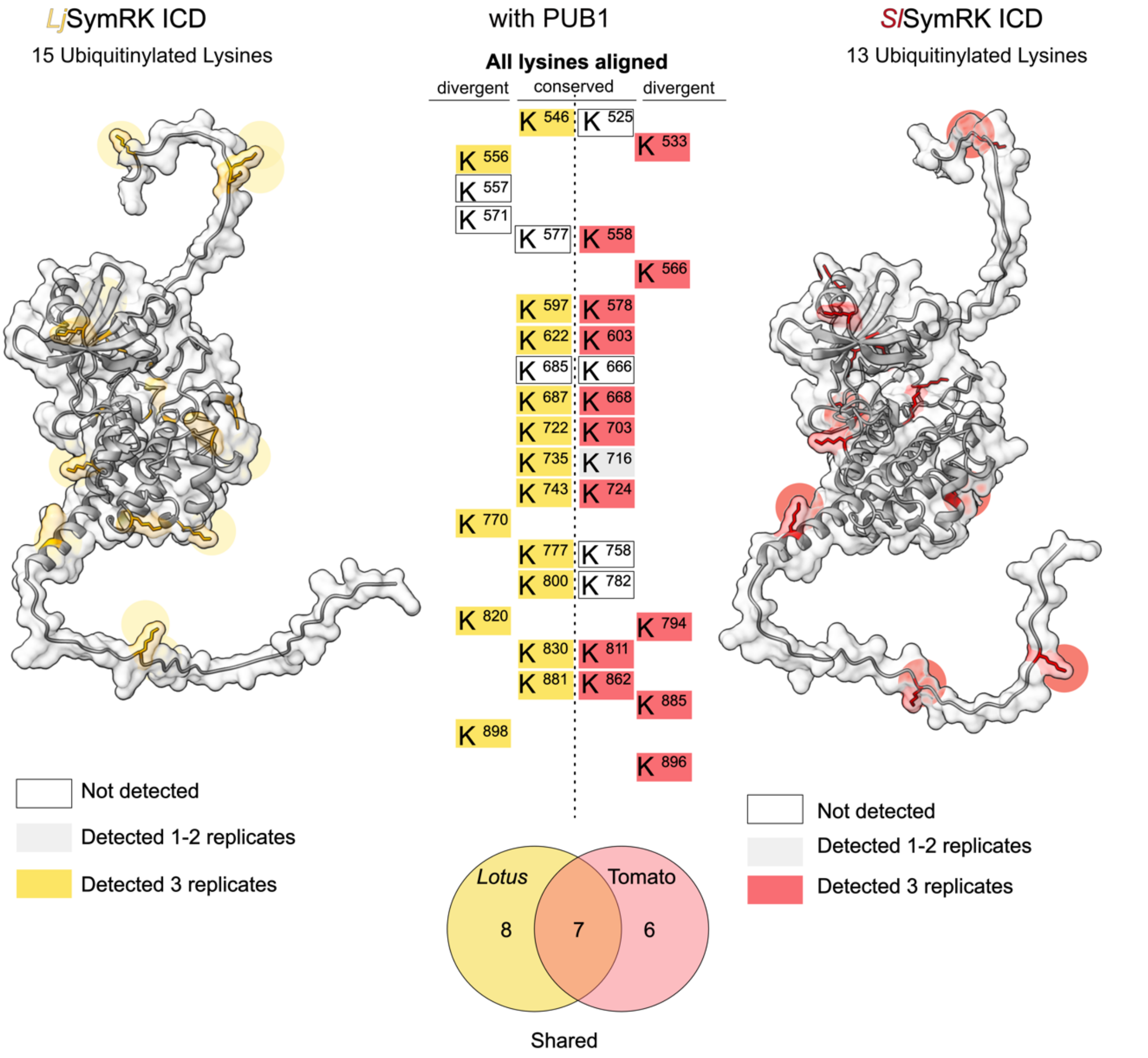


**Supplementary Figure 13. PUB1$^{\Delta N}$ ubiquitinylates *Lotus* and tomato SymRK ICDs at several lysine residues.**

The ICD of tomato (red) and *Lotus* (yellow) SymRK were expressed and purified from *E. coli*, subjected to *in vitro* ubiquitinylation assay with PUB1$^{\Delta N}$ followed by liquid-chromatography tandem mass spectrometry (LC-MS/MS) analysis to detect ubiquitinylated lysine residues. All *in vitro* ubiquitinylation assays were carried out three times. The sites were categorised as high confidence (all the replicates, yellow or red) vs low confidence (1-2/3 replicates, grey) and as not detected (white). Fifteen ubiquitinated lysine residues were identified on *Lotus* SymRK ICD with high confidence. On tomato SymRK, thirteen high-confidence lysine residues were detected. The identified sites were confirmed to be surface-exposed and mapped onto the corresponding AlphaFold 3-predicted structural models of SymRK ICDs from *Lotus* (left) and tomato (right). In both structural models, side chains of modified lysines are shown, and highlighted in red for tomato and in yellow for *Lotus* SymRK ICD. The list of modified lysines (K) and their respective amino acid identifiers is displayed in the central part of the Figure. Residues conserved in both SymRK variants based on amino acid sequence alignment are listed in the middle, while divergent residues appear at the periphery. The overlap between ubiquitinylated sites detected on tomato and *Lotus* SymRK ICD is depicted at the bottom of the Figure in a Venn diagram.

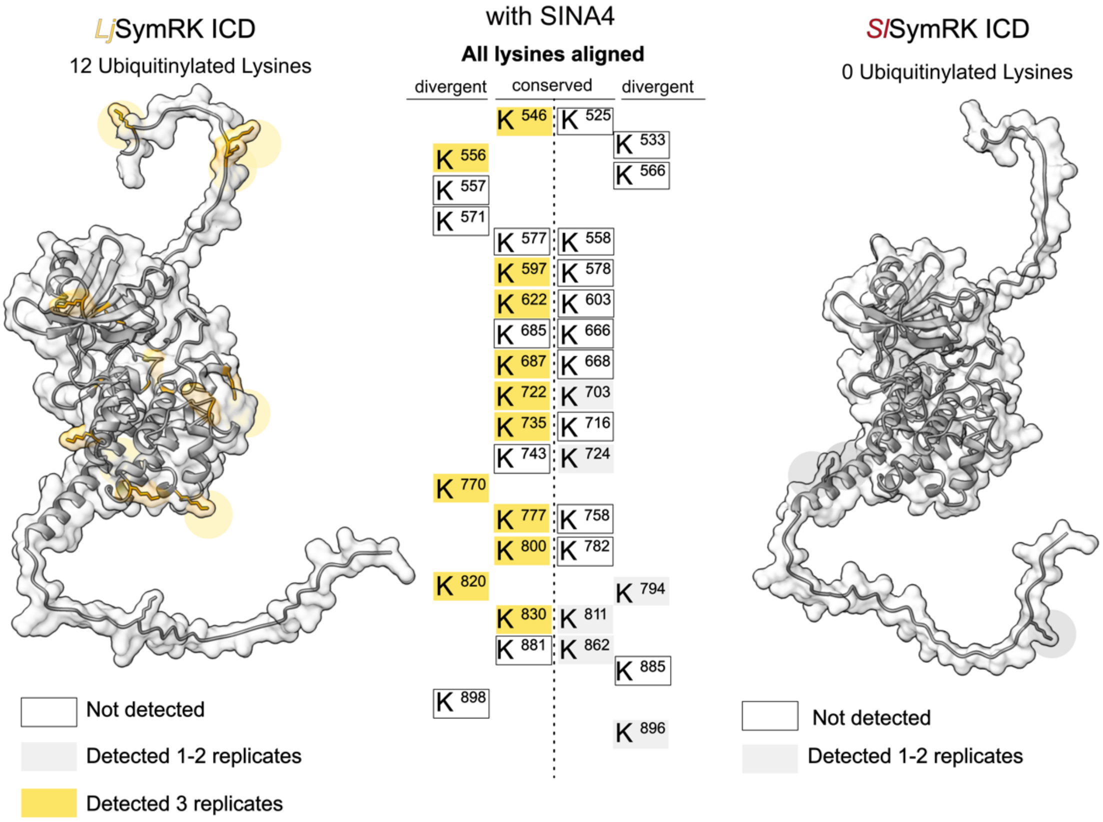


**Supplementary Figure 14. SINA4 ubiquitinylates the intracellular domain of *Lotus* SymRK at several lysine residues.**

The ICD of tomato (red) and *Lotus* (yellow) SymRK were expressed and purified from *E. coli*, subjected to *in vitro* ubiquitinylation assay with SINA4 followed by LC-MS/MS analysis to detect ubiquitinylated lysine residues. The sites were categorised as high confidence (3/3 replicates, yellow or red), low confidence (1-2/3 replicates, grey) and not detectable (white). Twelve ubiquitinylated lysine residues were identified on *Lotus* SymRK ICD in more than three replicates (highlighted in yellow). Ubiquitinylation was not detected with good reproducibility for tomato SymRK, supporting the observed lack of interaction between these proteins. The identified sites on *Lotus* SymRK ICD were confirmed to be surface-exposed and mapped onto the corresponding AlphaFold 3 predicted structural model. The list of all lysines (K) and their respective amino acid identifiers is displayed in the central part of the Figure. Residues conserved in both SymRK variants are located centrally, while divergent residues appear at the periphery, based on sequence alignment.

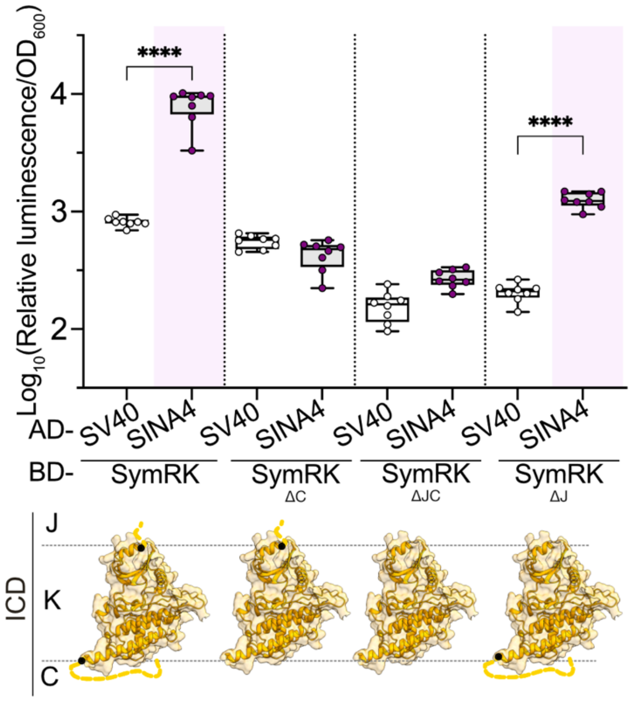


**Supplementary Figure 15. The unstructured C-terminal region of *Lj*SymRK is necessary for the yeast-two hybrid interaction with SINA4.**

Yeast two-hybrid interaction studies between SINA4 and different variants of SymRK ICD. The boxplots represent normalized luciferase signal measured for independent yeast colonies co-transformed with a construct encoding the SymRK ICD fused to the Gal4 Binding domain (BD) and SINA4 fused to the Gal 4 Activation Domain (AD), respectively. Simian-Virus 40 (SV40)-AD was used as a negative control. A series of *Lj*SymRK ICD deletion variants was tested that lack the juxtamembrane region (J), the unstructured C-tail (C) or both, as illustrated in the AlphaFold 3 models below the graph. Note that SymRK variants without the C-terminal region (SymRKΔC and SymRKΔJC) do not interact with SINA4. A yeast strain carrying an integrated *proGal2:Luciferase* reporter was used in the studies. One-way ANOVA followed by Tukey's multiple comparisons test was performed to assess significance. P-values < 0.05 are indicated with asterisks. J: juxtamembrane, K: core kinase, C: unstructured C-tail.
